# *Chd8* haploinsufficiency leads to molecular layer heterotopias and age-dependent cortical expansion

**DOI:** 10.64898/2026.02.28.708624

**Authors:** Felix A Kyere, Ian Curtin, Ziquan Wei, Cesar P Canales, Nicolas Seban, Lei Xing, Preslava P Todorova, Leticia Pérez-Sisqués, Maya Yilan Yin, Tao Wen, Roza M Vlasova, Dirgha Sheth, Jeffrey Bennett, Kejia Li, Nana Matoba, Carolyn M McCormick, Tala Farah, Oleh Krupa, Madison R Glass, Bonnie Taylor-Blake, Eric S McCoy, Tzu-Wen Winnie Wang, Qiuhong He, Mustafa Dere, Brooke R. D’Arcy, Liam T Davis, Veda Dayananda, Carla Escobar-Tomlienovich, Karthik Eswar, Maryam Moghul, Meghana Yeturi, Karen Huang, Micah Baldonado, Mihir Kaikini, David G Amaral, Debra L. Silver, David Borland, Hong Yi, Pablo Ariel, Yen-Yu Ian Shih, M Albert Basson, Laura C Andreae, Alex S Nord, Mark J Zylka, Guorong Wu, Jason L Stein

## Abstract

Mutations in the chromatin remodeler *CHD8* are associated with autism and macrocephaly. While mouse models of *Chd8* haploinsufficiency recapitulate brain overgrowth, the specific cellular mechanisms and developmental timing that lead to these anatomical abnormalities remain poorly understood. Here, we conducted 3D imaging of *Chd8*^V986*/+^ mouse brains using magnetic resonance imaging followed by tissue clearing and cellular resolution light-sheet microscopy across embryonic and postnatal developmental stages. We found that brain overgrowth occurs postnatally, driven by an increase in non-neuronal cells prior to volumetric expansion. Unexpectedly, we identified prevalent molecular layer heterotopias (MLH) within the frontal cortex of *Chd8*^V986*/+^ mice composed of neurons breaking through the pial surface during embryonic development and persisting throughout life. Increased incidence of MLH, previously identified in individuals with idiopathic autism, was replicated across three independent *Chd8^+/-^* mouse models and present across multiple genomic backgrounds, establishing aberrant neuronal migration as a core feature of *Chd8* haploinsufficiency.

## Introduction

Brain structural changes, including macrocephaly and cortical disorganization at the cellular level, have been observed in autistic individuals and mouse models harboring autism associated mutations^1–7^. However, the extent and cause of structural alterations in genetically defined forms of autism remains poorly studied.

Recent whole exome and whole genome sequencing efforts implicated hundreds of genes in ASD etiology^8–10^. Among the high-confidence genetic variants are *de novo* heterozygous loss-of-function mutations in the gene encoding Chromodomain helicase DNA-binding protein 8 (*CHD8*)^8,11,12^. Clinically, individuals with disruptive *CHD8* mutations often present with a distinct syndrome characterized by intellectual disability, dysmorphic facial features, gastrointestinal problems, and macrocephaly^11,13–15^. Macrocephaly is observed in a high percentage of patients with *CHD8* mutations (up to 85%). Mice carrying heterozygous mutations in *Chd8* display several ASD-related phenotypes, including increased brain weight or macrocephaly, mirroring clinical observations^16–21^.

CHD8 belongs to the chromodomain helicase DNA-binding protein family and functions as an ATP-dependent chromatin remodeler^22,23^. *Chd8* expression peaks during mid-gestation with highest levels observed in cortical progenitors and the marginal zone^24–26^. Disruption of *Chd8* impacts gene regulation^27^, progenitor proliferation^19,28^, and expression of genes involved with focal adhesion between cells^17,21^.

Despite the established link between *Chd8* haploinsufficiency and macrocephaly, the developmental trajectory of this overgrowth remains poorly defined. Specifically, it is unknown whether macrocephaly arises from a uniform, global increase in overall cell number, global increase in specific cell types, or if it is driven by focal disruptions in cortical architecture that traditional histology cannot resolve. Previous efforts to understand the cellular basis of *Chd8*-related macrocephaly focused on gross brain structural changes using MRI, histological imaging of 2D slices, or sequencing gene expression from individual cells, which lack spatial resolution to study individual cells, sample only a limited portion of the brain, or lack information on the location of those cells^16–21,26,29^.

Tissue clearing and rapid, high-resolution imaging methods have enabled intact whole-brain, cellular-resolution analysis to identify spatially resolved cell type specific impacts^30–32^. While one previous study has used tissue clearing to understand structural changes due to *Chd8* haploinsufficiency, the limited sample size, temporal, and spatial resolution prevent strong conclusions^33^. Here, we provide a comprehensive 3D brain structural analysis of a mouse line we generated that carries a pathogenic human *CHD8* stop codon mutation in exon 27 (*Chd8^V986*/+^*) through a combination of gross brain structural imaging using magnetic resonance imaging (MRI) and tissue clearing followed by light-sheet fluorescence microscopy (LSFM) to allow for whole-brain analysis at cellular resolution^17,30–33^.

Using our multi-modal approach across three developmental timepoints in mice (E18.5, P4 and P14), we define the onset of *Chd8^V986*/+^*-associated phenotypes. We report that macrocephaly is a postnatal event emerging between P4 and P14 driven by increases in non-neuronal cells as well as previously undescribed cortical heterotopias in the frontal cortex which are prevalent in embryonic cortex and persist throughout the lifespan. We also demonstrate that these heterotopias are replicated across different *Chd8* mutant mouse models and genetic backgrounds. This work identifies a direct link between autism-associated gene mutation and the emergence of cortical dysplasia.

### A multi-modal 3D pipeline for cellular-resolution analysis

To identify the brain regions and cell types impacted by *Chd8* haploinsufficiency across development, we established a multi-modal and 3D imaging and analysis pipeline (**Figure 1A**). Brains from *Chd8* heterozygous mice (*Chd8^V986*/+^*) on a verified C57BL/6J background (**Figure S1**) and their sex-matched wild-type littermates (WT) were collected at key neurodevelopmental time points: embryonic day 18.5 (E18.5), postnatal day 4 (P4), and postnatal day 14 (P14) yielding 118 total mice imaged (**Table S1**). Following initial *ex vivo* structural MRI to quantify brain volumes (at P4 and P14), intact brains or hemispheres underwent tissue clearing, were stained with a pan-nuclear marker (TO-PRO-3), and immunolabeled with cell-type and layer-specific markers (at E18.5, P4, and P14). Independent analysis of both hemispheres revealed no significant inter-hemispheric volume differences between *Chd8^V986*/+^* and WT littermate pairs (**Figure S2**). Consequently, the left hemisphere was arbitrarily selected for subsequent light-sheet imaging to optimize acquisition efficiency at E18.5 and P14. We processed hemispheres rather than full brains at these two ages to reduce imaging time and data sizes for different reasons: high resolution dual view imaging was needed at E18.5 when nuclei are densely packed despite the smaller volume of the brain and lower resolution but larger scanning volume was needed at P14 due to the larger size of the brain. Cleared and labeled brains (P4) or hemispheres (E18.5, P14) were then imaged in their entirety at cellular resolution using LSFM, enabling the generation of multi-channel, three-dimensional annotated datasets for detailed anatomical investigation.

**Figure 1:**
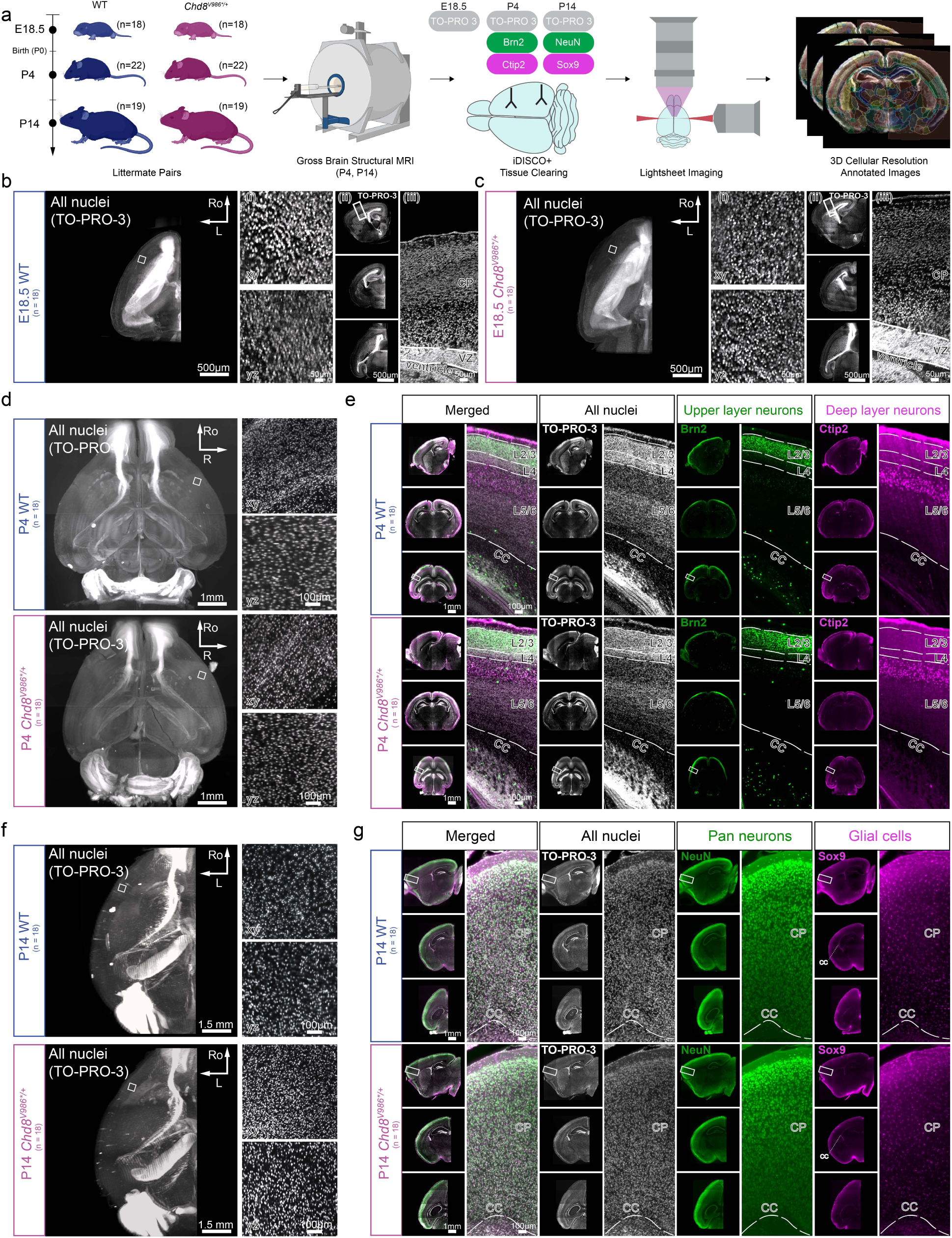
Experimental pipeline and representative images for characterizing the cellular basis of macrocephaly in *Chd8^V986*/+^* mice. a: Overview of the 3D imaging pipeline. Mice and MRI cartoons were generated with Biorender. b,c: Representative light sheet images of the left hemisphere from WT (b) and *Chd8^V986*/+^* (c) littermate pairs at E18.5 labeled with TO-PRO-3 (all nuclei). Left panel: 3D hemisphere rendering. (i): xy and yz sections from dual view deconvolved images showing single nuclei resolution. (ii): sagittal, coronal and axial sections. (iii): cortical plate (CP) and ventricular zone (VZ). Ro = Rostral; L = Left. d: 3D renderings of all nuclei of whole brain from WT and *Chd8^V986*/+^* littermate pairs at P4 labeled with TO-PRO-3 (all nuclei). Box insert represents xy and yz sections showing single nuclei resolution. Ro = Rostral; R = Right. e: Representative light sheet images of whole brain from WT and *Chd8^V986*/+^* littermate pairs at P4 labeled with TO-PRO-3 (all nuclei; grey) and upper layer neuronal marker (Brn2; green) and deep layer neuronal marker (Ctip2; magenta). Images include views of sagittal, coronal and axial sections and sections showing the layers of the isocortex and corpus callosum (CC). f: 3D renderings of all nuclei of whole brain from WT and *Chd8^V986*/+^*littermate pairs at P14 labeled with TO-PRO-3 (all nuclei). Box insert represents xy and yz sections showing single nuclei resolution. Ro = Rostral; L = Left. g: Representative light sheet images of left hemisphere from WT and *Chd8^V986*/+^* littermate pairs at P14 labeled with TO-PRO-3 (all nuclei), pan neuronal marker (NeuN) and a glial marker (Sox9). Images include views of sagittal, coronal and axial sections and sections showing the layers of the cortical plate (CP) and corpus callosum (CC).

Imaging of brains stained with the pan-nuclear marker provided a detailed overview of the cytoarchitecture at each developmental stage. Due to the dense packing of nuclei in the embryonic brain, we implemented a dual view light-sheet imaging approach by rotating the sample followed by joint deconvolution to obtain sufficient resolution to distinguish features in the image^34–36^ (**Figure S3**). At E18.5, both WT (**Figure 1B**) and *Chd8^V986*/+^* (**Figure 1C**) brains displayed stereotypical cell densities within the laminae of the cortical wall characteristic of late embryonic development. By P4 and P14, the continued growth, expansion, and cellular organization of the cortex and major brain structures were clearly visible in the 3D rendering of the brain from both genotypes (**Figure 1D,F**). This whole-brain visualization at cellular resolution provides the foundation for performing whole-brain cell counting, and identification and characterization of specific structural abnormalities.

To visualize the organization of distinct neuronal and glial populations, we utilized a panel of immunofluorescent markers that label specific cortical layers and cell types. At P4, labeling for Ctip2, a marker for deep-layer neurons (Layers Vb/VI), and Brn2, a marker for upper-layer neurons (Layers II/III), revealed the expected laminar patterns^37^ (**Figure 1E, Video S1**). At P14, we labeled NeuN^38^, a pan-neuronal marker, which densely labeled neurons throughout the isocortex and subcortical structures. In addition, labeling for Sox9^39^, a transcription factor expressed in a population of astrocytes and oligodendrocyte progenitors at P14^40^, identified these glial cells throughout the brain tissue, with higher density found in the corpus callosum as expected (**Figure 1G, Video S2**). The high resolution multi-modal dataset enables the identification of distinct expression patterns of these markers in WT brains confirming their utility for identifying and quantifying specific cell populations, and assessing the integrity of cortical lamination in *Chd8^V986*/+^*mice.

### Macrocephaly emerges postnatally between P4 and P14

We quantified the gross anatomical consequences of *Chd8* haploinsufficiency on brain volume at P4 and P14 through deformation of MR images to an age matched atlas (**Figure S4**)^41^. As expected, tissue clearing led to significant sample shrinkage, hence volumes were measured exclusively using MRI prior to tissue clearing (**Figure S5; Table S11**).

Representative MR images from both WT and *Chd8^V986*/+^* at P4 (**Figure 2A**) and P14 (**Figure 2C**) demonstrate sufficient resolution and contrast to clearly identify major anatomical structures, such as the isocortex and hippocampus. At P4, our volumetric analysis revealed no statistically significant differences between *Chd8^V986*/+^* and their WT littermates (**Figure 2B**). This was true for total brain volume (npairs=19) as well as for specific brain regions, including the isocortex and in both sexes.

**Figure 2:**
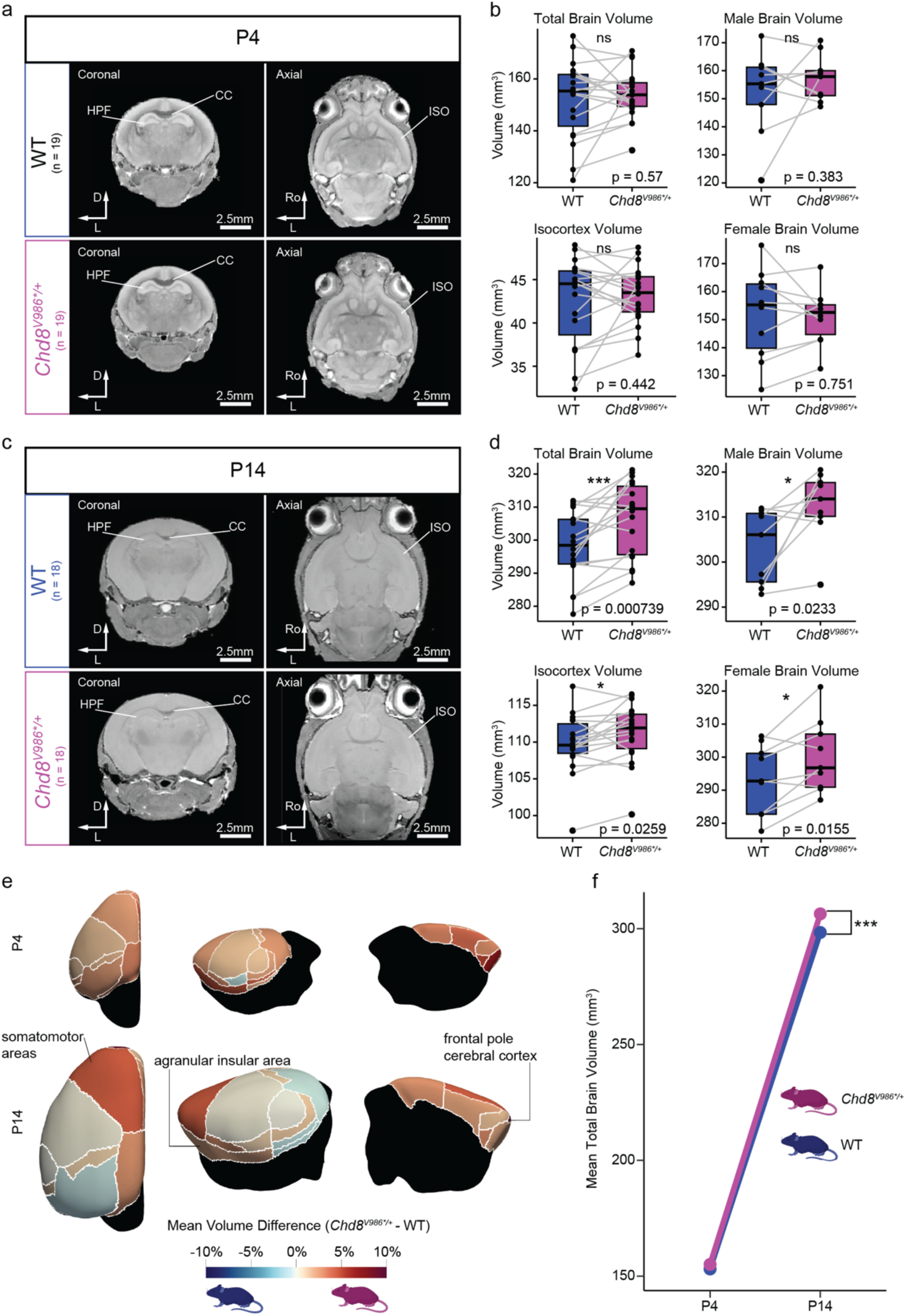
*Chd8^V986*/+^* macrocephaly is age dependent and region specific. a: Representative coronal and axial sections of structural MRI data at P4. ISO = Isocortex; HPF = Hippocampal formation; CC = Corpus Callosum; D=Dorsal; L=Left; Ro=Rostral b: Total brain (n_pairs_=19), male brain (n_pairs_=9), female brain (n_pairs_=10), and isocortex volume (n_pairs_=19) at P4 across genotypes. P-values are evaluated using a multiple linear regression accounting for sex-matched littermate pairs (equivalent to a paired t-test). Lines indicate sex matched littermate pairs. Boxplots show the first to third quartiles with whiskers representing 1.5 times the interquartile range. ns represents non-significant p-values. c: Representative coronal and axial sections of structural MRI data at P14. ISO = Isocortex; HPF = Hippocampal formation; CC = Corpus Callosum; D=Dorsal; L=Left; Ro=Rostral. d: Total brain (n_pairs_=18), male brain (n_pairs_=9), female brain (n_pairs_=9), and isocortex volume (n_pairs_=18) at P14 across genotypes. P-values are evaluated using a multiple linear regression accounting for sex-matched littermate pairs. Lines indicate sex matched littermate pairs. Boxplots show the first to third quartiles with whiskers representing 1.5 times the interquartile range. * represents p<0.05; *** represents p<0.001. e: Mean bilateral volume difference in isocortex regions across genotype at P4 (n_pairs_=19) and P14 (n_pairs_=18) evaluated through a multiple linear regression. Significant regions are labeled (FDR<0.1). f: Summary of the trajectory of brain overgrowth. *** denotes significant brain volume difference between at P14 (P-value from multiple linear regression, p = 0.000739; n_pairs_=18).

In contrast, a significant overgrowth was evident in the brains of *Chd8^V986*/+^* mice at P14 (**Figure 2D**). Total brain volume was significantly increased (npairs=18; 2.69% mean increase). This overgrowth was observed in both male and female sex-matched littermate pairs and was also seen in several specific brain structures including the isocortex. These findings demonstrate that the characteristic macrocephaly associated with *Chd8* mutations emerges during a postnatal developmental window between P4 and P14.

To further investigate the spatial and temporal dynamics of this brain overgrowth, we mapped the mean volume differences across various cortical subregions. While volume increases are subtle at P4, they become more pronounced by P14, with significant increases in average bilateral somatomotor, agranular insular area, and frontal pole cortical areas at this time point (**Figure 2E; Table S2;** FDR < 0.1). Examination of the developmental growth trajectory confirms the postnatal onset of macrocephaly. While no detectable difference in brain volumes were observed at P4, the *Chd8^V986*/+^* mice displayed an accelerated growth trend, resulting in an increased total brain volume compared to WT controls by P14 (**Figure 2F**).

### Increases in cell density precede cell number and volume expansion

To determine the cellular basis underlying the macrocephaly observed in *Chd8^V986*/+^* mutants, we performed whole-brain or hemisphere nuclear segmentation at P4 and P14, respectively. To segment nuclei, we developed a highly accurate extension of CellPose that we call nuclei instance segmentation (NIS) which stitches nuclei across patches and can be efficiently applied to whole mouse brains^42,43^ (**Figure S6**). In the left hemisphere of the WT isocortex, we detected around ∼6 million nuclei at P4 and ∼7.6 million nuclei at P14. These counts are consistent with existing studies quantifying cell types in the mouse brain^44^ (**Figure 3A,D; Table S3**; **Figure S7**). To investigate whether cellular count preceded volumetric changes observed at P14 (**Figure 2**), we compared nuclei number within the isocortex across genotypes at P4. We observed a non-significant trend toward higher cell count at P4 in *Chd8^V986*/+^* as compared to WT. The bilateral summed cell counts within isocortex regions were not significantly different across genotype at this age (**Figure 3B**). To account for variations in structural volume, we quantified cellular density alongside absolute cell counts. We observed significant increases in *Chd8^V986*/+^* global isocortex cell density at P4, with significant effects also detected in visual regions (**Figure 3C**; FDR<0.1). The increases in density preceded both higher global and regional isocortex cell counts and density at P14 (**Figure 3E,F**). Significant elevations in cell counts were found in the P14 auditory, visual, orbital and retrosplenial areas among others (**Figure 3E**). Mapping regional density differences revealed that while most regions showed modest changes, the visual areas displayed a significant increase in cell density (**Figure 3F, Table S4**). Together, this suggests that early postnatal alterations in cellular density precede subsequent changes in absolute brain volume.

**Figure 3:**
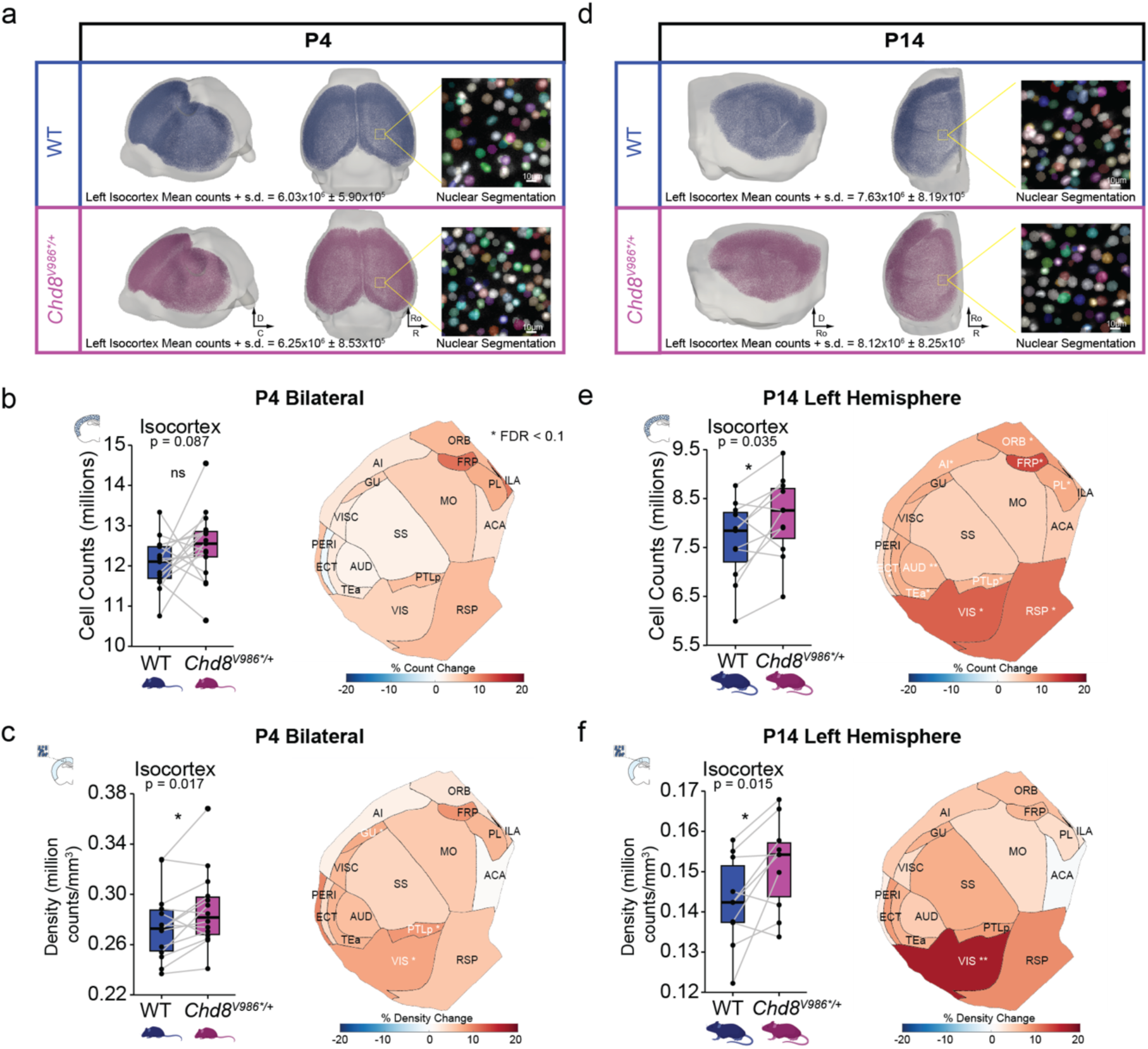
Increased cell count and density in *Chd8^V986*/+^*isocortex. a: Whole-brain or hemisphere nuclear segmentation at P4. Renderings of nuclei density in the isocortex from representative WT and *Chd8^V986*/+^*brains. Average and standard deviation of nuclei counts in each genotype are shown. Insets demonstrate the nuclear instance segmentations as colors over greyscale TO-PRO-3 images. b: Nuclei counts across genotypes in the isocortex and individual regions for summed bilateral effects at P4 (npairs = 17). Boxplots show the first to third quartiles with whiskers representing 1.5 times the interquartile range. P-value is evaluated using a multiple linear regression controlling for tissue loss and pair. ns represents non-significant p-value. c: Density of nuclei as in b (npairs = 14). P-value is evaluated using a multiple linear regression controlling for tissue loss and pair. In boxplot, * represents p<0.05. In flatmap, * represents FDR < 0.1. d: Whole-brain or hemisphere nuclear segmentation at P14. Renderings of nuclei density in the isocortex from representative WT and *Chd8^V986*/+^* brains. Average and standard deviation of nuclei counts in each genotype are shown. Insets demonstrate the nuclear instance segmentations as colors over greyscale TO-PRO-3 images. e: Nuclei counts across genotypes in the isocortex and individual regions for summed left hemisphere differences across genotypes at P14 (npairs = 11). Boxplots show the first to third quartiles with whiskers representing 1.5 times the interquartile range. P-value is evaluated using a multiple linear regression controlling for tissue loss and pair. In boxplot, * represents p<0.05. In flatmap, * FDR < 0.1, ** FDR < 0.01, *** FDR < 0.001. f: Density of nuclei as in e (npairs = 10). P-value is evaluated using a multiple linear regression controlling for tissue loss and pair. In boxplot, * represents p<0.05. In flatmap, * FDR < 0.1, ** FDR < 0.01, *** FDR < 0.001.

### Expansion of non-neuronal populations drives cortical overgrowth

To resolve the cell types driving the macrocephaly observed in *Chd8^V986*/+^* mutants, we performed whole-brain nuclear segmentation and cell-type classification at both P4, prior to macrocephaly, and P14, the time point when volumetric overgrowth is detected.

We first trained a multi-channel colocalization algorithm to classify individual nucleus instances within the P4 cortex based on expression of Brn2 and Ctip2 **(Figure 4A; Figure S6)**. Light-sheet images overlaid with segmentation masks confirmed high accuracy in identifying individual nuclei and their corresponding protein markers within the cortical architecture achieving high precision and recall scores for both Brn2 (0.85/0.9) and Ctip2 (0.91/0.89). Quantifying these populations across the entire P4 cortex and within the somatomotor area revealed no significant difference in Brn2 or Ctip2 neuronal cell counts, but a significant expansion of the Brn2-/Ctip2- populations (p = 0.004 whole cortex; FDR=0.027 somatomotor), which primarily consists of non-neuronal cells **(Figure 4B,C)**. To determine if this early expansion persists into later postnatal development, we evaluated nuclear counts at P14 using a similar multi-channel colocalization algorithm based on NeuN and Sox9, again achieving high precision and recall scores for both NeuN (0.95/0.95) and Sox9 (0.92/0.86) **(Figure 4D)**. Cell-type specific counts within the P14 left hemisphere cortex demonstrated that the established macrocephaly phenotype at this timepoint is driven by a significant expansion of NeuN-/Sox9- cells (p = 0.048), with no detectable changes in counts of NeuN+ neurons or Sox9+ glial cells **(Figure 4E,F; Table S5,S9)**.

**Figure 4:**
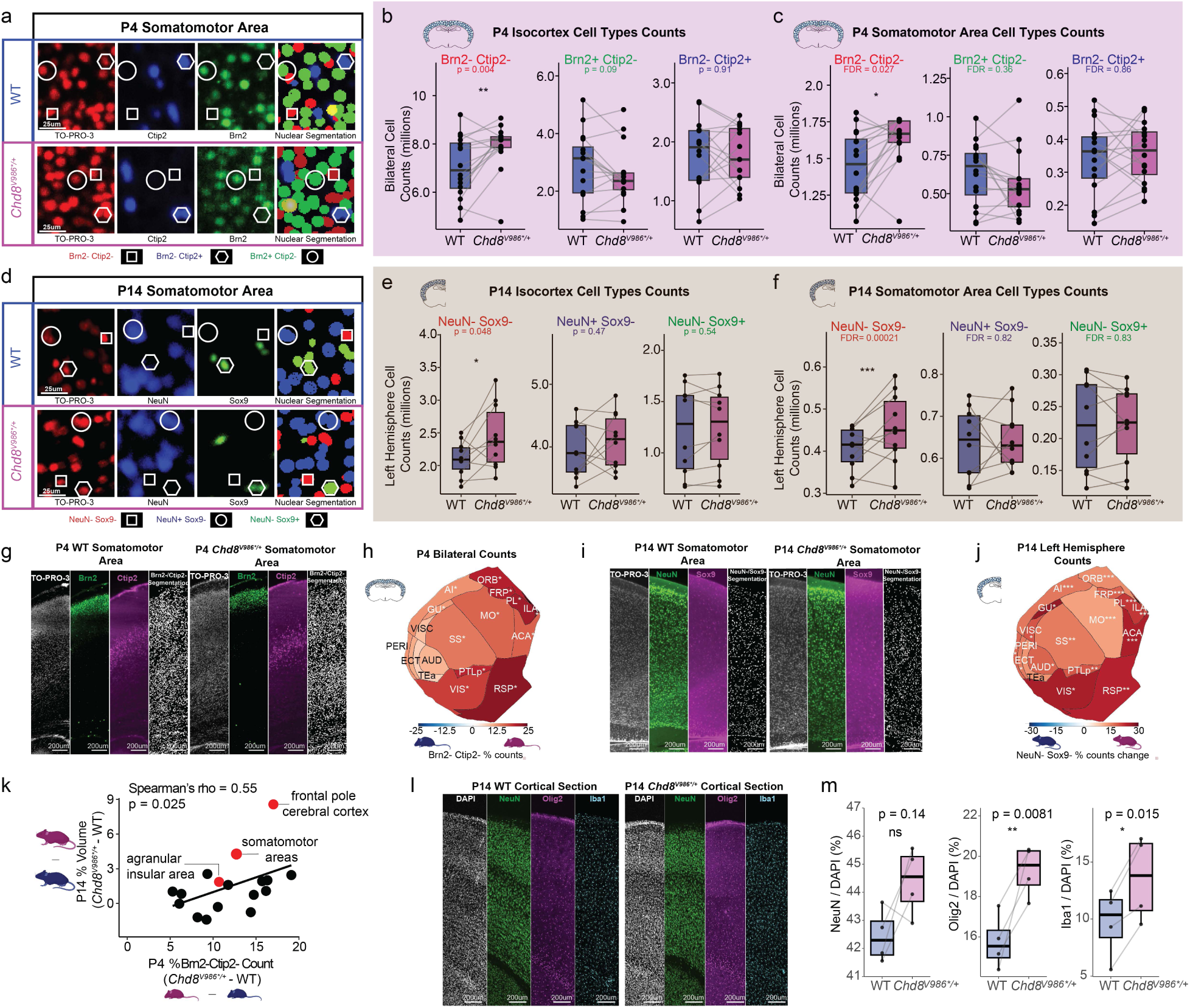
Expansion of non-neuronal populations drives *Chd8^V986*/+^* macrocephaly. a: Classification of nuclei within the somatomotor area of P4 WT and *Chd8^V986*/+^*mice. Sections show a nuclear marker (TO-PRO-3), the upper layer neuronal marker Brn2, and the lower layer neuronal marker Ctip2. The “Nuclear Segmentation” panel illustrates the cell-type classification into three populations: Brn2- /Ctip2- (red dot), Brn2-/Ctip2+ (blue dot), and Brn2+/Ctip2- (green dot). Brn2+/Ctip2+ populations are rare and not shown. b: Global quantification of cells in the P4 isocortex for Brn2-/Ctip2-, Brn2+/Ctip2- and Brn2-/Ctip2+ populations (npairs = 16). Lines indicate littermate pairs with P-value evaluated using a multiple linear regression controlling for pair and tissue loss. In boxplot, * represents p<0.05. c: Quantification of cells in the P4 somatomotor area for Brn2-/Ctip2-, Brn2+/Ctip2- and Brn2-/Ctip2+ populations (npairs = 16). Lines indicate littermate pairs with P-value evaluated using a multiple linear regression controlling for pair and tissue loss (* FDR < 0.1, ** FDR < 0.01, *** FDR < 0.001). d: Classification of nuclei within the somatomotor area of P14 WT and *Chd8^V986*/+^*mice. Sections show a nuclear marker (TO-PRO-3), the neuronal marker NeuN, and the glial marker Sox9. The “Nuclear Segmentation” panel illustrates the cell-type classification into three populations: NeuN-/Sox9- (red dot), NeuN+/Sox9- (blue dot), and Neun-/Sox9+ (green dot). NeuN+/Sox9+ populations are rare and not shown. e: Global quantification of cells in the P14 left hemisphere isocortex for NeuN-/Sox9-, NeuN+/Sox9-neuronal, and NeuN-/Sox9+ glial populations (n_pairs_ = 10). Lines indicate littermate pairs with P-value evaluated using a multiple linear regression controlling for pair and tissue loss. In boxplot, * represents p<0.05. f: Quantification of cells in the P14 somatomotor area for NeuN-/Sox9-, NeuN+/Sox9- and NeuN-/Sox9+ populations. Lines indicate littermate pairs with P-value evaluated using a multiple linear regression controlling for pair and tissue loss (* FDR < 0.1, ** FDR < 0.01, *** FDR < 0.001). g: Representative images for TO-PRO-3, Brn2, Ctip2 and NIS segmentation of Brn2-Ctip2- cells in the somatomotor area of WT (left) and Chd8^V986*/+^ (right) mice. h: Flattened isocortex heatmap of P4 Brn2-/Ctip2- count changes displaying the percentage change in Brn2-/Ctip2- counts between Chd8^V986*/+^ and WT littermate pairs across 17 isocortical regions (n_pairs_ = 9). P-value evaluated using a multiple linear regression controlling for pair and tissue loss. Significance is indicated by False Discovery Rate (* FDR < 0.1, ** FDR < 0.01, *** FDR < 0.001). i: Representative images for TO-PRO-3, NeuN, Sox9 and NIS segmentation of NeuN-/Sox9- cells in the somatomotor area of WT (left) and Chd8^V986*/+^ (right) mice. j: Flattened isocortex heatmap of P14 NeuN-/Sox9- count changes displaying the percentage change in NeuN-/Sox9- counts between Chd8^V986*/+^ and WT littermate pairs across 17 isocortical regions (n_pairs_ = 9). P-value evaluated using a multiple linear regression controlling for pair and tissue loss. Significance is indicated by False Discovery Rate (* FDR < 0.1, ** FDR < 0.01, *** FDR < 0.001). k: Scatter plot comparing the percentage change in Brn2-/Ctip2- cell counts at P4 (x-axis) with the percentage change in volume at P14 (y-axis) across various brain regions. Significance evaluated through Spearman’s rho correlation. * represents p value < 0.05. l: Representative confocal images for DAPI, NeuN, Olig2 and Iba1 in the cortex of WT (left) and Chd8^V986*/+^ (right) mice. Scale bar = 200µm. m: Quantification of NeuN+, Olig2+, and Iba1+ populations expressed as a percentage of total DAPI+ nuclei in the cortex (n_pairs_ = 4). Lines indicate littermate pairs. Statistical significance determined using a two-tailed paired t-test.

To visualize the structural distribution of these changing populations across the cortex, we projected these changes onto a flattened cortical heatmap at P4, showing that this non-neuronal expansion is widespread across the entire cortex **(Figure 4G,H)**. Similarly, mapping these cell population changes across the P14 cortex revealed widespread distribution and persistent expansion of the NeuN-/Sox9- nuclear population at this later stage **(Figure 4I,J)**.

We also explored changes in cell density, which control for differences in volume. At P4, we found a significant increase in the density of the Brn2-Ctip2- population across the entire isocortex (p=0.013), with clear density increases also detected within the somatomotor cortex (FDR=0.029; **Figure S10A,B,C**). By P14, the density of the NeuN-Sox9- population was similarly elevated across the global isocortex (p=0.0067) and within the somatomotor cortex (FDR=0.033; **Figure S10D,E,F**). Together, these density changes mirror our absolute count data and show that the non-neuronal cells are both expanded and more tightly packed together in the isocortex in *Chd8^V986*/+^*mutants.

We next asked whether the magnitude of early cellular expansion directly predicts downstream volumetric overgrowth. Spatial correlation analysis revealed that the regional percentage change of the Brn2-/Ctip2- at P4 is strongly and positively correlated with regional volume expansion at P14 (Spearman’s rho = 0.55, p = 0.025) **(Figure 4K).** Early cell-type changes at P4 precede the overall volumetric differences observed at P14, suggesting that localized cellular expansion is an early driver of the later brain volume overgrowth.

To resolve the molecular identities making up this expanding double-negative population, we performed high-resolution quantitative immunohistochemical imaging on P14 cortical sections **(Figure 4I)**. Quantitative analysis of these sections revealed that while the proportion of NeuN+ neurons relative to total DAPI+ cells is not significantly changed (p = 0.14), *Chd8^V986*/+^* mice display a significant increase in both Olig2+ oligodendrocyte lineage cells (p = 0.0081) and Iba1+ microglia (p = 0.015) **(Figure 4M)**. Collectively, these results demonstrate that *Chd8*-related macrocephaly is a result of increased early expansion of non-neuronal populations that drive postnatal cortical overgrowth.

### Early-embryonic formation of molecular layer heterotopias in frontal cortex

We examined the laminar structure of the cortex at the three key developmental timepoints: E18.5, P4, and P14. In the vast majority of WT control animals, the upper layers of the cortex displayed a well-organized laminar structure with sparse nuclei in Layer I (**Figure 5A, top row**). Unexpectedly, the cortices of many *Chd8^V986*/+^* littermates exhibited mushroom-shaped cell-dense incursions into Layer I that appeared to break beyond the pial surface at E18.5 (**Figure 5A, lower panels; Table S1; Videos S3,S4 ,S5**). The malformations were present at all time points sampled in a subset of *Chd8^V986*/+^*mice. Similar structures were previously named molecular layer heterotopias (MLH)^45,46^ and are likely caused by a breakage in the pial surface during embryonic development allowing over migration of neurons. Although not visible under the dissection scope at the early developmental timepoints, MLH can be observed in dissected brains in brightfield without tissue processing at 4 weeks (**Figure 5B**). To quantify the extent of this phenotype, we measured the presence, number, and volume of MLH in WT and *Chd8^V986*/+^* brains across the developmental timepoints. MLH were found in 28 (47% of total) of the *Chd8^V986*/+^* brains compared to 5 (8% of total) in their WT littermate pairs. Detection of MLH in WT mice is consistent with previous work which shows they are present in the C57BL/6J strain^45^. However, a higher prevalence of MLH were found at each age sampled (E18.5: p = 0.00772; P4: p = 0.0088; P14: p = 0.042) (**Figure 5C**). The presence of heterotopias may be underestimated by sampling only hemispheres at E18.5 and P14, as compared to full brains at P4. Heterotopias were found in both male and female mice, in pups derived from different sires and dams, and were not driven by unintended strain contamination during breeding (**Figure S1, S8**). We detected several animals with more than one MLH and there was an increase in the number of heterotopias per *Chd8^V986*/+^*animal at E18.5 (p = 0.00326), P4 (p = 0.00205), and P14 (p = 0.0195) (**Figure 5D**). A similar trend was observed in volumetric analysis. The total volume occupied by these heterotopic structures in the cortices of the mouse brains was significantly greater in *Chd8^V986*/+^* mice at E18.5 (p = 0.0044), P4 (p = 0.0027), and P14 (p=0.016) (**Figure 5E**). Interestingly, the MLH structures were found to be similar volumes across all timepoints sampled, which suggests that these structures were fully formed by E18.5 and the mechanism causing these malformations occurs at early embryonic stages (**Figure S8E**). Larger MLH were also visible as subtle signal intensities in MRI prior to tissue clearing from the same brains (**Figure S9**). This discovery prompted us to determine if cortical heterotopias were present in older mice. We found that 80% (n=4 of 5) of 4-week old *Chd8^V986*/+^* mice had heterotopias that were visible under a dissecting microscope relative to 0% (n=0 of 3) in age-matched WT animals (**Figure 5B**), suggesting that the percentage of *Chd8^V986*/+^* animals with heterotopias is stable across the lifespan. Together, these data demonstrate that the *Chd8^V986*/+^* mutation leads to embryonic formation of MLH that persist in the cortex.

**Figure 5:**
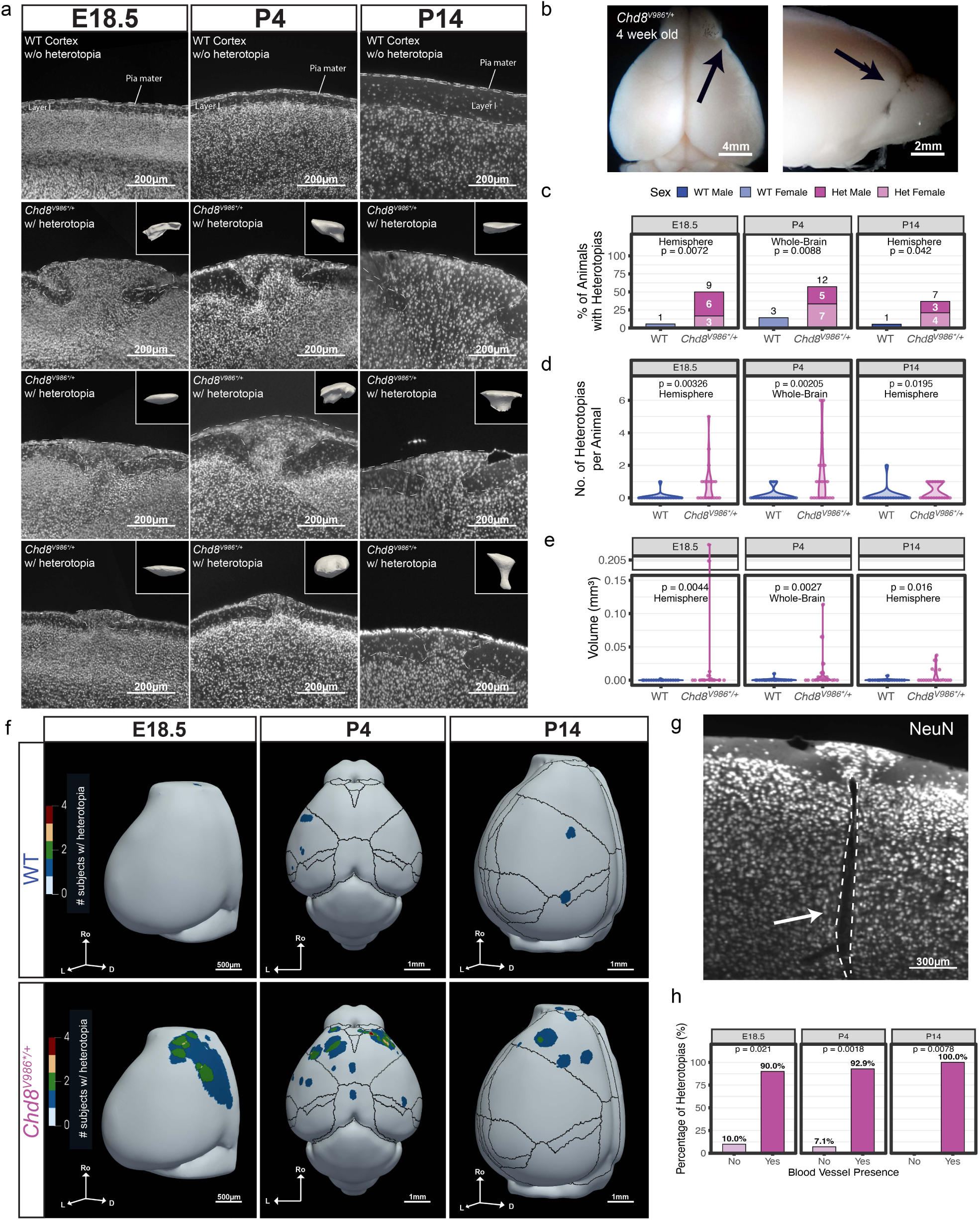
Molecular layer heterotopias in *Chd8^V986*/+^* cortex. a: Representative coronal sections of the cortex at embryonic (E18.5) and postnatal (P4 and P14) stages. Wild type (WT) animals generally have organized laminar structure (top row), while *Chd8^V986*/+^* exhibit disruption of cortical layers by mushroom-shaped heterotopias (bottom rows). Heterotopias are outlined in dashed lines. Insets display 3D volumetric renderings of the segmented heterotopias. b: Macroscopic view of a 4-week-old *Chd8^V986*/+^* brain. Arrows indicate heterotopias visible on the cortical surface. c: Quantification of heterotopia prevalence across genotype at each age. Bar graphs show the percentage of animals where heterotopias were detected, stratified by sex (blue = WT male, light blue = WT female; grey = Het male, pink = Het Female). P-values indicate significant differences between genotypes evaluated by a two-sided Fisher’s Exact Test (E18.5 npairs = 18, P4 npairs = 22, P14 npairs = 19). d: The number of heterotopias per animal quantified across genotype at each age, displayed as violin plots. Differences were evaluated through a two-sided Wilcoxon Rank-Sum test (E18.5 npairs = 18, P4 npairs = 22, P14 npairs = 19). e: Volumetric comparisons of the heterotopias across genotypes at each age, displayed as violin plots. Differences were evaluated through a two-sided Wilcoxon Rank-Sum test (E18.5 npairs = 18, P4 npairs = 22, P14 npairs = 19). f: Three-dimensional spatial reconstruction of heterotopia distribution. Heatmaps map the frequency of heterotopias onto a consensus brain shape, revealing a preferential clustering in the frontal cortex of *Chd8^V986*/+^*mice compared to WT controls. D=dorsal; L=lateral; Ro=rostral. g: Representative image showing a physical association between heterotopias and the brain vasculature. A blood vessel (outlined in dashed white) extends from the ventricle toward the base of the heterotopia (arrows). h: Quantification of the association between heterotopias and blood vessels across development. Bar graphs show the percentage of heterotopias containing blood vessels at E18.5, P4, and P14. Differences were evaluated through a two-sided Binomial test (n = 33).

To determine if these cortical malformations had a specific spatial distribution, we generated three-dimensional reconstructions of the reference mouse brain at each age and mapped the locations of all identified heterotopias. In WT animals, very few heterotopias were detected at any age and were never in the most frontal regions (**Figure 5F, top row**). However, MLH in *Chd8^V986*/+^* mice were strongly enriched in the frontal cortex (**Figure 5F, bottom row**). This clustering in the frontal cortex suggests a region-specific cortical malformation caused by *Chd8* haploinsufficiency. These cortical heterotopias may have been missed in prior *Chd8* mutant mouse studies because they occupy a small volume and are not found in consistent locations, so slice immunohistochemistry would not likely have included heterotopias and lower resolution MRI is barely sufficient to resolve these small structures. In contrast, tissue cleared brain imaging has both high spatial resolution and whole brain sampling enabling detection of MLH.

We noted a frequent association between the MLH and tube-like structures assumed to be brain vasculature. In almost all MLH, blood vessels were seen extending from the ventricle toward the base of the heterotopic structures (**Figure 5G**). We quantified the presence of blood vessels within heterotopias and show that this enrichment is highly significant across developmental stages suggesting a stable neuro-vascular interaction within these malformations (**Figure 5H**).

### The MLH phenotype is replicated across distinct *Chd8* mutant lines and genetic backgrounds

To establish that the MLH phenotype is a consistent feature of *Chd8* heterozygosity rather than specific to a single allele or genetic background, we examined cortical architecture across three independent experimental models. First, we evaluated an independent *Chd8* mutant line maintained on a C57BL/6J background harboring a 5-bp deletion in Exon 5 (*Chd8^5bpdel/+^*) likely leading to frameshift mutations and nonsense mediated decay^19,47^ at P42 using serial Nissl staining across the entire cortex **(Figure 6A,B)**. These mice exhibited mushroom-shaped heterotopias that were morphologically indistinguishable compared to the malformations characterized in our primary model. We identified a significant increase in the percentage of *Chd8^5bpdel/+^* animals presenting with MLH compared to wild-type littermates (p = 0.018) **(Figure 6C)**. MLH were predominantly in the frontal cortex in this model as well. Second, we assessed another independent line harboring an Exon 3 deletion (*Chd8^ΔEx3/+^*) on the C57BL/6J background^48^ at P49 using immunohistochemistry where serial slices were acquired from only the frontal cortex **(Figure 6D,E)**. NeuN immunolabeling highlighted aberrant neuronal clusters in the upper layers of the cortex mirroring the aberrant cellular organization found in *Chd8^V986*/+^*mice and demonstrating a significantly increased prevalence of MLH compared to wild-type controls (p = 0.041) **(Figure 6F)**. Finally, because C57BL/6J mice are known to exhibit a baseline susceptibility to sporadic MLH^45^, we investigated whether the phenotype persists on a hybrid genetic background resistant to heterotopia formation. Previous studies detected no MLH in wild-type C57BL/6J x DBA/2J F1 hybrid crosses^49^. When we crossed *Chd8^V986*/+^*C57BL/6J males to DBA/2J females and examined F1 B6-DBA offspring at P30 **(Figure 6G,H)**, we observed NeuN+ heterotopias in the upper cortical layers of multiple *Chd8^V986*/+^*F1 B6-DBA mutants trending toward higher prevalence as compared to littermate controls (p = 0.074) **(Figure 6I)**. This replication across multiple mutant mouse lines, ages, and strains demonstrates that the heterotopias are not specific to given mutations in *Chd8* but are characteristic of multiple types of loss of function mutations, strengthening evidence that *Chd8* haploinsufficiency causes MLH.

**Figure 6:**
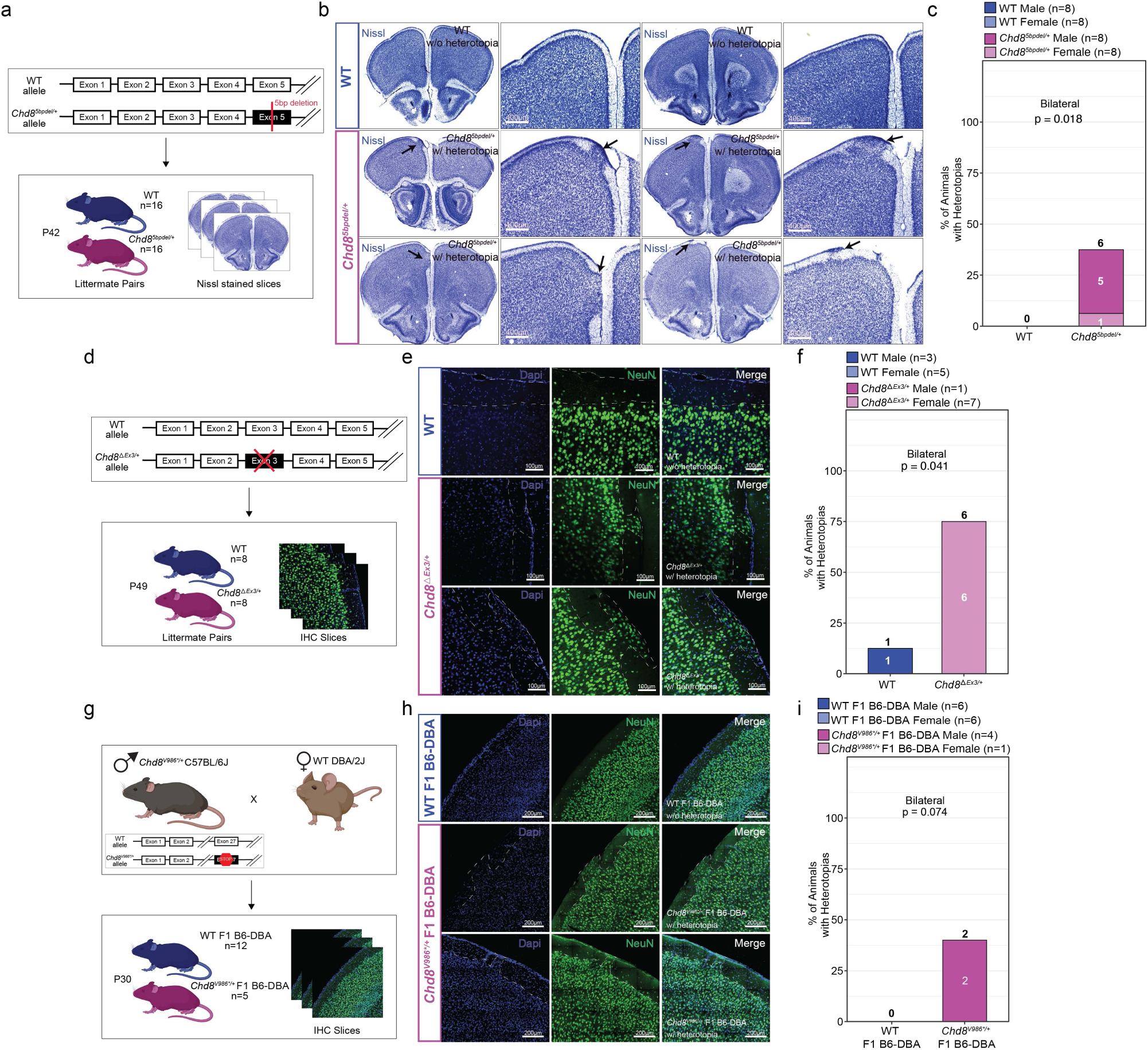
Molecular Layer Heterotopias (MLH) are replicated across independent *Chd8* mutant lines and genetic backgrounds a: Schematic overview of the targeting strategy disrupting Exon 5 and the experimental workflow utilizing P42 littermate pairs (n = 16 WT, n = 16 *Chd8^5bpdel/+^*). b: Representative coronal brain sections stained with Nissl. Arrows indicate MLH within the upper cortical layers of *Chd8^5bpdel/+^* mice. Scale bars = 400μm. c: Quantification of the percentage of animals presenting with MLH. Numbers within or above bars denote individual animal counts stratified by sex. P-values calculated by Fisher’s exact test. d: Schematic overview of the Exon 3 deletion allele and immunohistochemistry experimental design (n = 8 WT, n = 8 *Chd8^ΔEx3/+^)*. e: Representative confocal images of cortical sections immunolabeled for DAPI, NeuN, and merged channels. Scale bars = 100μm. f: Quantification of MLH penetrance. P-values calculated by Fisher’s exact test. g: Breeding scheme crossing male *Chd8^V986*/+^* C57BL/6J mice to female WT DBA/2J mice to generate F1 hybrid offspring (n = 12 WT, n = 5 *Chd8^V986*/+^* F1 B6-DBA). h: Confocal images of P30 F1 hybrid cortical sections stained for DAPI and NeuN. Scale bars = 200μm. i: Quantification of MLH penetrance in F1 hybrids demonstrating a trend toward increased MLH incidence in mutants. P-values calculated by Fisher’s exact test.

### Cell type composition of molecular layer heterotopias

Following the identification of MLH in *Chd8^V986*/+^* mice, we next sought to characterize their cellular composition to understand their origins and impact. Using the cell type specific markers in tissue cleared images, we found contrasting cell type composition in small versus large heterotopias. In large heterotopias, we observed a mixed population of cells, with abundant expression of both Ctip2 and Brn2 (**Figure 7A, top row**). This indicates that these larger structures are composed of a disorganized combination of neurons originating from both deep and superficial cortical layers, consistent with a broad disruption of neuronal positioning throughout corticogenesis. The composition was different in smaller heterotopias, which were almost exclusively composed of Brn2-positive, upper-layer neurons, with a notable absence of Ctip2-positive, deep-layer neurons (**Figure 7A, bottom row**). This finding suggests that the neuronal composition of the heterotopia may vary depending on the timing of the pial breakage, as more neurons can migrate past an earlier pial boundary breakage leading to both upper and layer neurons forming a larger heterotopia.

**Figure 7:**
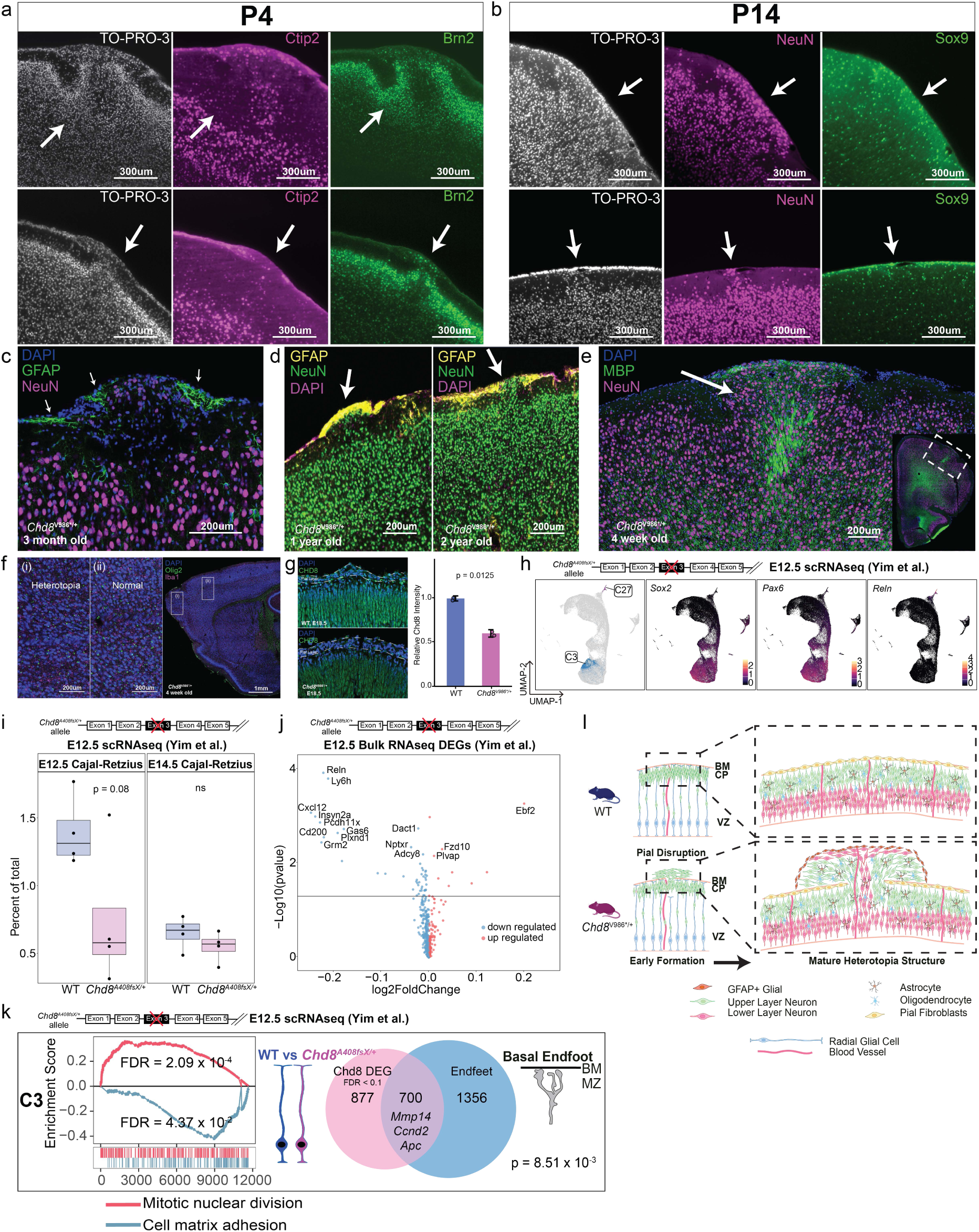
Cellular composition and proposed mechanism of *Chd8^+/-^* molecular layer heterotopias. a: Immunohistochemical characterization of neuronal identity within heterotopias at P4 in *Chd8^V986*/+^*. Top row: Large heterotopias contain a mixed population of early-born deep-layer neurons (Ctip2+, magenta) and later-born upper-layer neurons (Brn2+, green). Bottom row: Smaller heterotopias are composed almost exclusively of upper layer neurons (Brn2+). TO-PRO-3 (white) marks nuclei. b: Analysis of neuronal and glial composition of heterotopias at P14 in *Chd8^V986*/+^*. Heterotopias are densely packed with neurons (NeuN+, magenta), while the distribution of glial cells (Sox9+, green) appears unperturbed and comparable to surrounding tissue. c: Immunolabeling for Gfap (green) and NeuN (magenta) shows the accumulation of astrocytes at the cortical surface site above the heterotopia. d: Representative images of long-term structural analysis at 1 year and 2 years showing the presence of an astrocyte cap (Gfap+, yellow) above the heterotopia. e: Staining for Myelin Basic Protein (MBP, green) and NeuN reveals myelinated tracts projecting into or from the heterotopic clusters. f: Representative images comparing the densities of oligodendrocytes (Olig2+, green) and microglia (Iba1+, magenta in a heterotopia (i) to the adjacent normal cortex (ii). g: CHD8 immunolabeling at pial boundary at E18.5 in WT and *Chd8^V986*/+^*embryos and quantification of relative CHD8 intensity. Pial layers are outlined with dashed lines. Differences were evaluated with a two-sided Welch’s Two-Sample T-test (n*_Chd8V986*/+_* = 2; n_WT_ = 2). h: UMAP visualization of scRNA seq in developing mouse cortex from a previous study^26^. Cells are colored by expression of radial glial markers *Sox2* and *Pax6* (middle and right, respectively highly expressed in radial glial cluster C3), and *Reln* expression in Cajal-Retzius cells (C27). i: Box plots showing the percentage of Cajal-Retzius cells (C27) as a proportion of total recovered cells from scRNA-seq datasets at E12.5 (left) and E14.5 (right). Wild-type (WT) controls are shown in blue and *Chd8^A408fsX/+^* mutants are shown in magenta. P-values are calculated using a two-tailed unpaired Student’s t-test. j: Volcano plot displaying differentially expressed genes (DEGs) subset to Cajal-Retzius marker genes from E12.5 bulk RNA-seq data (blue dots indicate downregulated genes; red dots indicate upregulated genes) k: Gene set enrichment analysis (GSEA) for differentially expressed genes between *Chd8^A408fsX/+^*and WT within a selected radial glial cluster (C3) at E12.5 (left panel). GSEA score curves for mitotic nuclear division (pink) and Cell matrix adhesion (blue) are shown. X axis indicates the position of genes contributing to the enrichment along the ranked list. Venn diagram showing DEGs in C3 overlapped with radial glia endfeet localized genes^54^ (right panel), with selected overlapping genes listed. P-value represents significance evaluated from a two-sided Fisher’s Exact Test. BM, basement membrane; MZ, marginal zone. l: Proposed model for the development of molecular layer heterotopias and macrocephaly. The schematic illustrates the timeline from early embryonic pial breach to adult cellular composition and glial cap.

In both large and small heterotopias at P14, the irregular clusters of cells were densely positive for NeuN, confirming that the structures consist mainly of misplaced neurons (**Figure 7B, top and bottom rows**). NeuN does not label Cajal-Retzius cells residing in cortical Layer I^50^, and the aberrant NeuN+ clusters observed in the heterotopias indicate the presence of heterotopic cells. In contrast, the spatial pattern of Sox9-positive glial cells appeared largely unperturbed. The glial cells were distributed evenly throughout the cortex and did not show any obvious accumulation within or around the neuronal heterotopias, suggesting that the primary defect lies in neuronal migration rather than a broader disorganization of both neuronal and glial cell types.

We examined the long-term consequences of these developmental malformations in adult *Chd8^V986*/+^* mice at 3 months, 1 year, and 2 years of age. At 3 months, GFAP+ astrocytes began to accumulate at the heterotopic breach site, indicating that astrocytes were migrating to this location (**Figure 7C**). Labeling for NeuN and GFAP at 1 year and 2 years of age revealed that the mushroom-shaped heterotopias were persistently capped by a dense layer of GFAP-positive cells (**Figure 7D**).

To determine the cellular maturity and integration of the heterotopias in juvenile mice (4 weeks), we performed immunohistochemistry for neuronal and glial markers. Strikingly, labeling myelin basic protein (MBP) revealed dense, aberrant myelinated tracts projecting directly into or from the heterotopic clusters (**Figure 7E**), suggesting that these ectopic neurons are receiving and/or projecting axonal connections and are not isolated from the wider cortical circuit. We also labeled oligodendrocytes (OLIG2) and microglia (IBA1) and viewed no difference in the density of these cells in an individual section with a heterotopia as compared to surrounding tissue (**Figure 7F**).

To understand a potential mechanism whereby *Chd8* haploinsufficiency leads to MLH, we immunolabeled CHD8 in embryonic brains. CHD8 is highly expressed in cells near the pial surface at E18.5, as has been detailed previously^25,26,51^. As expected, a significant reduction in CHD8 protein intensity at the pial boundary was observed in *Chd8^V986*/+^* embryos compared to WT littermates (p = 0.0125) (**Figure 7G**). We speculate that the reduction of CHD8 expression may weaken the pial boundary during early embryonic development, a mechanism which has been described for heterotopia formation caused by other mutations^52^.

To test this hypothesis, we re-analyzed scRNA-seq data from the developing cortex at E12.5 from embryos harboring an exon 3 heterozygous mutation^26^, *Chd8^A408fsX/+^*. UMAP visualization confirmed the expression of canonical Cajal-Retzius neuron marker *Reln* (**Figure 7H**). Quantifying cell type proportions across early cortical development revealed a trending decrease in the percentage of *Reln*-positive Cajal-Retzius cells at E12.5 in *Chd8^A408fsX/+^* compared to wild-type controls, an effect that normalized by E14.5 (**Figure 7I**). To provide additional evidence, we examined bulk RNA-seq data from the E12.5 from the same study. We observed strong down-regulation of Cajal-Retzius cell markers, including *Reln*, *Ly6h*, and *Cxcl12*, in *Chd8^A408fsX/+^* embryos consistent with a reduction in Cajal-Retzius cells (**Figure 7J**). Cajal-Retzius cells secrete Reelin to stop neuronal migration, so an early embryonic reduction in Cajal-Retzius cells in *Chd8^+/-^*embryos may lead to migration past the pial boundary and heterotopia formation.

A complementary mechanism leading to heterotopia formation is inconsistent tiling of radial glial endfeet that creates holes in the cortex allowing neuronal migration beyond the pial boundary^53^. To study this mechanism, we utilized a cluster expressing radial glial markers (*Pax6* and *Sox2*) from *Chd8^A408fsX/+^* E12.5 embryos^26^. Gene set enrichment analysis (GSEA) identified a significant upregulation of genes associated with mitotic nuclear division across these progenitor clusters while cell-matrix adhesion genes were significantly downregulated in *Chd8^A408fsX/+^* embryos (**Figure 7K**). Interestingly, radial glial DEGs significantly overlapped with transcripts localized to the radial glial basal endfeet^54^, including those involved in formation of the basement membrane and endfoot adhesion (*Mmp14, Ccnd2, Apc*)^55–58^. We next sought to replicate this finding in an independent, E14.5 snRNA-seq dataset utilizing the *Chd8^5bpdel/+^* mouse line (**Figure S11A**). Genes dysregulated by *Chd8* haploinsufficiency in radial glial cells again demonstrated a significant transcriptomic overlap with the same set of transcripts highly expressed in radial glial endfeet, including overlapping targets *Ccnd2* and *Apc,* as well as *Fabp7* (**Figure S11B**). Because *Chd8* is known to bind directly to the promoters or regulatory regions of *Mmp14*, *Ccnd2*, *Fabp7* and *Apc* to modulate their expression^59^, our findings point to a mechanism where *Chd8* mutations destabilize downstream cascades essential for structural radial glial endfeet adhesion. In humans, mutations in *CCND2* disrupt cortical architecture and manifest as polymicrogyria^60,61^, while *APC* is vital for establishing radial glial polarity, with its loss driving severe neuronal migration dysregulation^62,63^. Together, our results suggest that *Chd8* haploinsufficiency compromises the cell adhesion, migration, and structural signaling programs, leading to neuronal migration beyond the pial boundary and heterotopia formation with the eventual formation of an astrocyte cap at the location of the pial breach (**Figure 7L**).

## Discussion

In this study, we utilized a high-resolution 3D imaging pipeline to provide a comprehensive spatio-temporal map of brain abnormalities in the *Chd8^V986*/+^* mouse model. Our multi-modal approach revealed two developmentally distinct brain structural phenotypes caused by *Chd8* haploinsufficiency: (1) the embryonic establishment of neuronal molecular layer heterotopias (MLH) in frontal cortex which are persistent throughout life, and (2) postnatal macrocephaly driven by increased cell generation of oligodendrocyte and microglial populations. Given the different developmental trajectories (embryonic establishment of heterotopias but postnatal establishment of macrocephaly) and cell type composition (neuronal composition of heterotopias but glial cells driving macrocephaly) of these two phenotypes, we expect that *Chd8* haploinsufficiency impacts two separate aspects of brain development.

Many mouse models have been generated to study *Chd8* haploinsufficiency, as well as emerging non-human primate models, almost all of which replicated macrocephaly observed in humans^19,20,48,64–67^. However, the temporal emergence of macrocephaly, the regional localization, and the cellular composition have remained largely unknown. Previous studies using Nissl staining in cortical slices from *Chd8*^+/-^ mice demonstrated that macrocephaly was not driven by neuronal populations^47^. Consistent with this, our data demonstrate that this overgrowth is a postnatal event emerging between P4 and P14, driven predominantly by an increase in oligodendrocytes and microglia. Specifically, whole-brain nuclear segmentation reveals that the significant elevations in cell counts across the isocortex are attributed to a Sox9-/NeuN-population, suggesting that neither neurons (NeuN+ cells) nor astrocytes or oligodendrocyte progenitor cells (Sox9+ cells) are substantially contributing to cortical expansion. Macrocephaly, a defining clinical hallmark of *CHD8* syndrome, appears to be driven by an atypical expansion of glial lineages. Quantitative immunofluorescence of the *Chd8^V986*/+^* cortex at P14, a period characterized by peak microglial density^68,69^ and active oligodendrocyte maturation^70^, revealed significant increases in both Iba1+ microglia and Olig2+ oligodendrocyte lineage cells. These findings align with the established role of *CHD8* as a master regulator of oligodendrocyte maturation^71^, recent observations of increased oligodendrocyte density in *Chd8^+/-^* primates^67^, and the demonstration that microglial-specific *Chd8* depletion impairs oligodendrocyte differentiation^72^. Notably, the observed glial overgrowth occurs between P4 and P14, coinciding with the peak of cortical myelination and microglial expansion^68,69,73,74^. Our data suggest that a dysregulation of the glial rather than neuronal overproduction, acts as the primary driver of *Chd8* haploinsufficiency induced macrocephaly.

We unexpectedly found that the neurodevelopmental pathology of *Chd8* mutations involves an alteration in neuronal localization, rather than solely a late-stage expansion of cell numbers. This phenotype was likely missed in previous studies of *Chd8*^+/-^ mice because heterotopias are incompletely penetrant, and not found in identical locations such that slice histology may not sample the heterotopia, and can be smaller than the resolution used for whole brain approaches like MRI. In fact, we observed heterotopias in archival immunohistochemical slices from *Chd8^V986*/+^*mice that were missed in our previous characterizations of this mouse^17^. The identification of heterotopias in a mouse modeling a mutation associated with a psychiatric disorder argues for the wider use of whole brain cellular resolution imaging in both mouse and human to determine whether similar small structural features are more broadly associated with risk for psychiatric disorders^75^. MLH were obvious in whole brain cellular images because of the high contrast of dense nuclei in the otherwise cell-sparse Layer I. Other types of heterotopias may be detectable if appropriate labels for the affected cell types are used. Finally, our analysis focused largely on the isocortex but structural changes may occur in other regions as well, which are able to be quantified through this whole brain imaging approach.

We found that MLH are a definitive consequence of *Chd8* haploinsufficiency and not an artifact of a single mouse line or a specific targeting strategy. By evaluating independent *Chd8* mutant lines targeting Exon 3 and Exon 5, we demonstrated that the significantly increased prevalence of MLH is reproducible across multiple heterozygous loss of function variants. Despite variation in mutation sites and the ages of the cohorts samples, these independent models exhibited mushroom-shaped malformations that were morphologically indistinguishable to those on our *Chd8^V986*/+^* line. We also demonstrated that the MLH phenotype is consistent across other genetic backgrounds by cross-breeding *Chd8^V986*/+^*males with WT DBA/2J females and observing an increased trend of MLH incidences in the mutant mice. This hybrid strain model shows that the MLH are present even on a genetic background resistant to sporadic MLH formation.

Molecular layer heterotopias are likely caused by damage to the pial surface during embryonic development, allowing migrating neurons to break through this barrier. The pial surface acts as a physical barrier, preventing migrating neurons from exiting the brain^52,76^. Molecular layer heterotopias can reproducibly be generated in mice and rats by physically damaging the pial surface with a micropipette during embryonic development^77–79^. Given the high expression of *Chd8* in the most basal cortical laminae during mid-gestation^25,26,51^, a reduction in Cajal-Retzius cells that normally stop over-migration of neurons, the altered expression of radial glial endfeet genes^54^ , and the relationship between heterotopia size and neuronal composition related to the timing of cortical development, we propose a model where *Chd8* haploinsufficiency decreases the number of reelin expressing cells and weakens the radial glial endfeet adhesion, allowing neurons to over-migrate into the subarachnoid space, as has been observed in other models^80–82^. This pial breach represents a permanent structural alteration. These ectopic neurons survive into adulthood, are capped by astrocytes, supplied with nutrients by vascularization, and become integrated into cortical function by receiving or projecting dense myelinated axonal connections. The spatial clustering of MLH within the frontal cortex is particularly interesting given the central role of this region in executive function and social cognition and their impairment in individuals with *CHD8* mutations^83–85^. These focal heterotopias represent potential “hotspots” of circuit dysfunction that may occur independently of global brain volume changes.

Molecular layer heterotopias, polymicrogyria, and other malformations of cortical development have been identified in post-mortem brains from individuals with autism, dyslexia, focal epilepsy, and other neurodevelopmental disorders, suggesting functional significance in human patients as well^5,86–97^. Cortical heterotopias have not yet been detected in MRI from patients harboring loss of function *CHD8* mutations, though only a small number of case studies or qualitative radiological reports are currently available and are complicated by low resolution and motion artifacts^98,99^. Given the distinctive cytoarchitectural and vascular abnormalities within these heterotopias (e.g., aberrant white matter bundles and abnormal vasculature), it may be possible to determine whether heterotopias are present in humans using MR imaging techniques optimized to detect these features^100,101^.

By uncovering the dual nature of *Chd8^V9^*^86*\*/+*^ pathology, we move closer to understanding the complex neuroanatomical landscape of autism. The precise molecular mechanism by which a chromatin remodeler alters pial boundary integrity may be elucidated using preparations that study gene expression specifically in radial glial endfeet in *Chd8^V986*/+^*mice or spatial transcriptomics^54^. Future work may also detail the extent to which the number, size and location of the heterotopias are associated with behavioral severities in *Chd8^V986*/+^*mice, similar to previous work which induced cortical disorganization using maternal immune activation^6^. Additionally the connectivity and activity of the heterotopic neurons should be evaluated through anterograde and retrograde labeling and electrophysiology^102^.

## Methods Animals

*Chd8* heterozygous mice (*Chd8^V986*/+^*) were bred by mating male *Chd8^V986*/+^*, previously generated through CRISPR genetic editing^17^, with female wildtype mice (Jackson laboratory). All mice were on the C57BL/6J strain. The mice were housed in a 12 hours:12 hours light-dark cycle with ad libitum access to food (Teklad 2020X, supplied by Envigo, Huntingdon, UK) and water. Dam and sire age, litter size and plug dam weight were recorded for each breeding event. To determine the sex and *Chd8* genotype, genomic DNA was extracted from tail clippings by Proteinase K enzyme digestion. *Chd8* genotyping was performed via PCR amplification of the genomic DNA using the following primers: (F) 5′ GCTAAGACAGAAATCTGATCTATTACCAGTAGA and (R) 5′ GGTCTTGAGATCCCCAAAATCCTTAA. This was followed by digestion with *Mbol* restriction enzymes to differentiate between the wild-type (WT; 227-bp product) and *Chd8*^V986*/+^ (149-bp and 78-bp products) alleles. Sex was determined through PCR amplification of the Y-chromosome-specific *Sry* gene. Presence or absence of the *Sry* band was visualized against a 1 kb DNA ladder to confirm male or female identity. Eight samples used for subsequent tissue clearing and imaging were additionally genotyped for genetic quality control with the MiniMUGA array^103^ and analyzed with the most updated pipeline as described previously^104,105^. This cohort included balanced sex distribution (4 males, 4 females), genotype distribution (nWT = 4 and n*Chd8V986*/+*= 4), and represented mice both with (n = 2) and without (n = 6) heterotopias. These samples were confirmed to be coisogenic mice carrying the *Chd8^V986*/+^* in pure C57BL/6J genetic background (**Figure S1**). We randomly selected sex-matched littermate pair pups at each of three time points, using one sex-matched pair per litter to reduce data dependency^106^. Timed matings were performed by pairing two WT adult females with one *Chd8^V986*/+^* male in the late afternoon. We checked dams for vaginal plugs no later than 12:00pm the following day and successful matings were then designated E0.5. For E18.5 experiments, pregnant dams were euthanized and embryos were harvested. P4 and P14 litters were harvested 4 and 14 days respectively after birth (P0). Each pup was marked with a unique ID and experimenters were blinded to the genotypes during downstream experiments. The animal protocols used were all approved by the Institutional Animal Care and Use Committee at the University of North Carolina at Chapel Hill.

### Sample Size Calculation

To ensure the study was adequately powered to detect neuroanatomical differences, a power analysis was conducted. Based on previous brain weight data^17^, we anticipated an increase in total brain volume in *Chd8* mutant mice compared to wild-type controls with effect size Cohen’s d=0.7. With the significance level set at 0.05 and the desired power at 80%, a sample size of *n* = 18 littermate pairs per time point was needed to detect significant effects. As a result, a total cohort of 108 brains was generated, along with 8 extra samples at P4 and 2 at P14, balanced by genotype, sex and developmental timepoint, providing sufficient sensitivity to detect increased brain volume (**Table S1**).

### MR Sample Preparation and Imaging

To evaluate brain structure at P4 and P14, Magnetic Resonance Imaging (MRI) was performed on 22 and 18 *Chd8*^V986*/+^ mice at P4 and P14 respectively along with their paired littermate controls (80 mice total), as described previously^30,107^. We did not conduct MRI imaging at E18.5 because the low resolution of MRI and small size of the brain prevents annotation of structures. Sex-matched littermate pairs were prepared in batches together and imaged in succession without knowledge of genotype during imaging. The mice underwent transcardial perfusion using PBS to wash out the blood followed by PBS + 4% paraformaldehyde (PFA) solution, which contained a 20% gadolinium-based MRI contrast agent (Prohance, Bracco Diagnostics, A9576, 0270-1111-03). The entire head of each pup was carefully decapitated at the base of the skull where the occipital bone meets the first cervical vertebra post-fixation, with the skin being carefully removed to leave the skull undamaged. The skulls containing the brain were weighed for each sample. The samples were then subjected to drop-fixation in a solution of PBS + 4% PFA for 24 hours at 4°C to ensure thorough fixation. Following this, the samples were immersed in PBS + 3% gadolinium solution for 23 days at 4°C. The samples were then placed securely in a 5mL syringe filled with Fomblin Y (Specialty Fluids Company, YL-VAC-25-6) solution and imaged using a Bruker 9.4T/30 cm horizontal-bore, animal MRI system. A 15 mm volume coil and a spin-echo-based sequence were used with the following parameters per sample: Spatial resolution: 60 µm x 60 µm x 60 µm; total scan duration of 7 hours 12 minutes; echo time (TE) of 6.83 ms; repetition time (TR) of 40 ms; excitation/refocusing flip angles of 90/180 degrees; image size of 166 x 168 x 209 voxels; Field of view (FOV) of 9.9 mm x 10.1 mm x 12.4 mm; and a bandwidth of 100,000 kHz. All 80 samples were batch processed in pairs (*Chd8*^V986*/+^ and WT sex-matched littermate controls). After MR imaging, samples were washed with PBS and stored in PBS + 1% Sodium Azide at 4°C until processed for tissue clearing and lightsheet imaging. Samples were stored for no more than one month.

### Tissue Clearing and Immunolabeling

Following MRI, the same intact brains with skull attached (22 *Chd8*^V986*/+^ and 22 WT littermate controls at P4; 18 *Chd8*^V986*/+^ and 18 WT littermate controls at P4) and the E18.5 samples (18 *Chd8*^V986*/+^ and 18 WT littermate controls) were tissue cleared with the iDISCO+ tissue clearing and immunolabeling protocol^30,31,108^. First, the skulls were carefully removed, the brain samples washed with PBS and then weighed. Subsequently, the samples underwent a dehydration process using a methanol gradient (Fisher, A412SK), a pretreatment with 66% dichloromethane (Sigma-Aldrich, 270997)/methanol and 5% H2O2 (Sigma-Aldrich, H1009)/methanol solution. This was then followed by rehydration, membrane permeabilization (20% dimethyl-sulfoxide (Fisher, BP2311); 1.6% Triton X-100 (Sigma-Aldrich, T8787); 23 mg/mL Glycine (Sigma-Aldrich, G7126), and blocking with 6% goat serum (Abcam, ab7481). The samples were then incubated for 4 days at 37°C in PTwH buffer solution (PBS; 0.5% Tween-20 (Fisher, BP337); 10 mg/L Heparin (Sigma-Aldrich, H3393) containing rabbit anti-Brn2 (Cell Signaling Technology, 12137, 1:100) and rat anti-Ctip2 (Abcam, ab18465, 1:400) for P4 brains, and 6 days with guinea pig anti-NeuN (Millipore-Sigma, ABN90P, 1:50) and rabbit anti-Sox9 (Millipore-Sigma, AB5535, 1:100) for P14. After a 2-day wash period with PTwH, the P4 samples were incubated with PTwH containing goat anti-rat Alexa Fluor 568 (Thermo Fisher, A11077, 1:200), and goat anti-rabbit Alexa Fluor 790 (Thermo Fisher, A11369, 1:50) for an additional 4 days at 37°C, and the P14 samples with donkey anti-guinea pig Alexa Fluor 594 (Jackson ImmunoResearch, 706-585-148, 1:50) and goat anti-rabbit Alexa Fluor 790 (Thermo Fisher, A11369, 1:100) for 6 days. All samples were incubated in TO-PRO-3 (Thermo Fisher, T3605, 1:400) for 4 days for E18.5 and P4, and 6 days for P14. Samples were then washed for 2 days with PTwH, followed by a second dehydration using a graded methanol series, a 3-hour incubation in 66% dichloromethane/methanol and a 30 minute incubation in 100% dichloromethane. The P4 and P14 samples were then stored in dibenzyl ether solution (RI = 1.56, Sigma-Aldrich, 108014) at room temperature. The E18.5 samples were stored in Ethyl Cinnamate (Millipore-Sigma, 8002380250). All 118 samples were batch processed in littermate pairs. Due to iDISCO+ and imaging-related errors, such as aberrant sample movements during image-capture, several MR-imaged samples were not properly imaged on the light sheet microscope and were omitted from this section of the study. All subsequent cellular resolution analysis and statistics were performed on this reduced dataset (**Table S1**). All intact samples were imaged on the light sheet microscope within one week of completing tissue-clearing and immunolabeling.

### Lightsheet Imaging

All postnatal iDISCO+ tissue clearing and immunolabeling samples were whole mount imaged using Ultramicroscope II light-sheet (LaVision Biotec) equipped with MVPLAPO 2X/0.5 NA objective with a corrected dipping cap (Olympus) , sCMOS camera (Andor), and ImSpector control software. E18.5 images were acquired using the Ultramicroscope Blaze (Miltenyi Biotec) equipped with a 12X/0.53 NA objective with a corrected dipping cap, sCMOS camera and Imspector control software. All samples were imaged as previously described^30,31,109^. Briefly, postnatal images were acquired at 0.75 µm x 0.75 µm x 4 µm/voxel with a single light sheet from the right, numerical aperture (NA) of 0.108 and magnification of 4x on the zoom body. E18.5 images were acquired at 0.22 µm x 0.22 µm x 1 µm/voxel with a single light sheet from the right, light sheet NA of 0.163 and magnification of 2.5X on the zoom body. To maintain axial resolution across the image width in postnatal brains, we employed dynamic horizontal focusing with the contrast-enhanced feature in ImSpector. The number of steps (typically 11 for all brains) utilized was recommended through the ImSpector software and dependent on the laser wavelength. To maintain axial resolution for E18.5 brains on the Blaze microscope, we utilized the “Light Speed” acquisition mode and enabled the “Fast Tiling Scan” feature with a scaling factor of 2.0. The P4 tissue cleared brains were positioned ventral-side down to allow for axial plane acquisition. The camera was positioned dorsally, perpendicular to a single light sheet facing the lateral side of the right hemisphere. As E18.5 and P14 samples were single hemispheres, these were positioned medial-side down on the cut edge to allow for sagittal plane acquisition, with the camera positioned laterally and the single light sheet entering the axial plane. The E18.5 samples were imaged a second time to enable isotropic resolution through a deconvolution and fusion framework as described below. Here, the brains were positioned with the caudal side down, the camera positioned rostrally and the single light sheet entering the sagittal plane. We employed a sequential approach for individual channels across a grid of 5 x 4 tile positions. To ensure comprehensive coverage and continuity, we maintained a 20% overlap between adjacent tiles at P4, 15% at P14 and 10% at E18.5. The excitation laser frequencies used were 785 nm (Brn2 in P4 and Sox9 in P14), 647 nm (TO-PRO-3 in all samples) and 561 nm (Ctip2 in P4 and NeuN in P14). For each tile, we collected stacks from sex-matched Chd8^V986*/+^ samples and their corresponding littermate controls. The z-dimension of these stacks typically ranged 4500 µm - 5000 µm, depending on the brain size. The duration of imaging sessions per sample at P4 and P14, for all three channels, varied between 13 to 15 hours and for 3 hours per E18.5 sample. This rigorous approach ensured the generation of nuclear-resolution intact-brain data sets for our subsequent analyses.

### Dual-View Volumetric Deconvolution and Isotropic Fusion

We employed a deep learning–based denoising and deconvolution framework^36^ to restore dual-view volumetric images, using a cascaded two-stage architecture consisting of a denoising module followed by a deconvolution module. The method is self-supervised and does not require clean or ground-truth images for training. During training, each view was processed independently using only the experimentally acquired noisy volume. Two training samples were generated by adding independent noise realizations, producing image pairs that share identical underlying structural content but differ in their noise components. One image was used as the network input, while the other served as the training target. The denoising module was trained using a fidelity-based loss that enforces consistency between the denoised prediction and the independently re-corrupted target. Because the noise realizations in the input and target are statistically independent, the network is forced to learn the shared structural signal rather than reproducing noise, thereby preventing trivial identity mapping. The denoised output was subsequently passed to a second-stage deconvolution network that explicitly incorporates the optical image formation model. The deconvolution loss included a degradation-consistency term, in which the deconvolved prediction was forward-blurred with a known point-spread function (PSF) and compared against the denoised image, together with a regularization term to stabilize high-frequency recovery and suppress noise amplification. At inference time, each noisy volume was processed in a single forward pass through the trained denoising and deconvolution modules, yielding a restored volume for each view. The reconstructed volumes from the two views were then transformed into a common coordinate system and combined using a weighted fusion strategy. The fusion weights were designed to exploit complementary information from the orthogonal viewing directions, emphasizing regions with higher local confidence while reducing residual anisotropic artifacts. This view-wise reconstruction followed by weighted fusion produced a final isotropic volume with improved contrast, reduced noise, and enhanced structural fidelity.

### Registration of Atlas to MR images

The Developmental Mouse Brain Templates provide, for each developmental stage, both developmental common coordinate framework (DevCCF) annotations and the Allen Common Coordinate Framework version 3 (CCFv3) annotation^41^. For each subject, age-matched P4 and P14 templates were nonlinearly registered to the individual skull-stripped MRI brain image in native space using a two-stage registration framework. First, an affine registration of the template MRI (ADC contrast) to the individual brain was performed using BRAINSFit (version 4.4.0). Registration was initialized using image moments (center of mass and principal axes), followed by rigid and affine registration. Similarity metrics were estimated using random sampling of 1,000,000 voxels to ensure robust and stable parameter estimation. Second, the resulting affine transform was used to initialize a nonlinear registration using ANTs (version 2.2.0;^110^). Nonlinear alignment was performed using the Symmetric Normalization (SyN) model with a gradient step size of 0.125. Image similarity was optimized using mutual information with two histogram bins, across a multiresolution scheme comprising 100, 75, and 50 iterations at coarse, intermediate, and fine resolution levels, respectively. Gaussian regularization was applied to both the deformation field (smoothing = 1) and the update field (smoothing = 0.5) to promote smooth and anatomically plausible deformations. The resulting ANTs affine transform was converted to an ITK-compatible format using ITKTransformTools (version 1.2.1). The combined affine transform and nonlinear deformation field were then applied to the Allen CCFv3 annotation using ResampleScalarVectorDWIVolume (version 0.1), resampling the labels into each animal’s native space with nearest-neighbor interpolation to preserve discrete regional identities. All resulting label maps were visually inspected to assess registration accuracy and anatomical plausibility.

### Registration of Atlas to Light-Sheet Images

Population-based anatomical templates for P4 and P14 were constructed from LSFM images using the ANTs multivariate template construction framework^111^. Skull-stripped individual density maps were iteratively aligned and averaged in three-dimensional space to generate unbiased, study-specific group templates. Template construction proceeded through four iterative refinement cycles. In each cycle, individual images were nonlinearly registered to the current template estimate using the Symmetric Normalization (SyN) model, gradient step size of 0.05, and the resulting deformation fields were used to update the template. All images were rigidly aligned prior to nonlinear refinement to minimize initial pose differences. Computation was parallelized across four CPU cores. The final study-specific templates represent the average anatomical structure of the P4 and P14 cohorts in a common coordinate space and exhibited reduced noise and improved tissue contrast. The corresponding age-matched LSFM templates (P4 and P14) from the Developmental Mouse Brain Templates^41^ were then nonlinearly warped to the study-specific templates. The CCFv3 annotation was propagated into the study-specific template space following the same registration and label-resampling procedures described in the Segmentation of MRI images section. From the study-specific template space, CCFv3 annotations were subsequently propagated to each animal’s native space using the inverse transformations obtained during template construction. Finally, the forward deformation fields computed during study-specific atlas building were used to transform heterotopia masks from individual native space into the study-specific template space for group-level analyses. Individual LSFM images lacked the cerebellum and olfactory bulbs, and P14 LSFM data were acquired from the left hemisphere only. To ensure accurate and unbiased registration, the corresponding P4 and P14 templates from the Developmental Mouse Brain Templates were manually masked (in ITK-SNAP v4.0.2^112^) to match the acquired anatomy prior to all registration procedures. This approach minimized bias introduced by partial or asymmetric image coverage by constraining registration to anatomically corresponding regions across subjects and templates. For the P14 study-specific template, limited manual refinement of the propagated CCFv3 annotation was performed using the 3D Slicer version 5.8.1^113^ “Landmark Registration” module to correct residual misalignments in cortical and subcortical structures. To mitigate the impact of physical defects on the NIS counting process, tissue cracks and dissection errors were manually annotated slice-by-slice in ITK-SNAP (v4.0.2) (**Figure S7D**). These regions were subsequently controlled for using linear models during downstream analyses.

### Brain Sections and Immunofluorescence Staining of *Chd8^V986*/+^* brains

Brains of embryonic mice (E18.5), one and two year old mice were dissected and drop-fixed in 4% PFA/PBS for 24 hours, rinsed with PBS and incubated in 30% sucrose/PBS at 4°C for 48 h and embedded in Epredia M-1 Embedding Matrix. Brains were sectioned using a Epredia™ CryoStar™ NX50 Cryostat at 50µm/section. P30 mice were perfused with 4% PFA/PBS and brains were dissected and post-fixed in 4% PFA overnight. P30 brains were embedded in 4% low-melting-point agarose/PBS and sectioned using a Leica VT1200 vibratome at 50 µm/section. Both embryonic and P30 brain sections were collected in PBS and stored at 4°C before immunofluorescence staining. Immunofluorescence staining was performed as described previously^31^. The following primary antibodies were used: goat anti-GFAP (Abcam, ab53554; 1:1000), guinea pig anti-NEUN (Millipore, ABN90P; 1:1,000), mouse anti-MBP (R&D Systems, MAB42282; 1:1000), rabbit anti-CHD8 (Bethyl, A301-225A; 1:500), rabbit anti-OLIG2 (Millipore, AB9610; 1:2,000), goat anti-OLIG2 (R&D Systems, AF2418; 1:1,000), rabbit anti-IBA1 (Wako, 019-19741; 1:1,000), and rat anti-CTIP2 (Abcam, ab18465; 1:1,000). The following secondary antibodies utilized include donkey anti-rabbit IgG Alexa Fluor 568 (ThermoFisher Scientific, A10042; 1:1,000), donkey anti-rat IgG Cy3 (Jackson ImmunoResearch, 712-165-153; 1:1,000), donkey anti-rat IgG Alexa 647 (Jackson ImmunoResearch, 712-605-153; 1:1,000), goat anti-mouse IgG1 Alexa Fluor 488 (ThermoFisher Scientific, A21121; 1:1,000), donkey anti-guinea pig IgG Alexa Fluor 647 (Jackson ImmunoResearch, 706-605-148; 1:1,000) or Alexa Fluor 488 (Jackson ImmunoResearch, 706-545-148; 1:1,000), donkey anti-goat IgG Cy3 (Jackson ImmunoResearch, 705-165-147; 1:1,000), and donkey anti-goat IgG Alexa Fluor 488 (Jackson ImmunoResearch, 705-545-147; 1:1,000). Nuclei were labeled by DAPI staining. Sections were mounted onto Fisherbrand Superfrost/Plus slides with an anti-fading Polyvinyl alcohol mounting medium with DABCO (Sigma-Aldrich, 10981). Heterotopia images were acquired using a Zeiss LSM 710 laser scanning confocal microscope equipped with three fluorescence spectral detectors. Excitation lasers used were: 405 nm for DAPI, 488nm for Alexa Fluor 488, 543 nm for Alexa Fluor 568 and Cy3, and 633 nm for Alexa Fluor 647. Emission filters used were: 406/545 for DAPI and Alexa Fluor 488, 551/632 for Alexa Fluor 568 and Cy3, and 637/756 for Alexa Fluor 647. Beam splitters used were MBS 488/543/633 and MBS 405. Original images were obtained using a 20X Plan Apochromat objective (NA: 0.8; WD: 0.55 mm; **Figure 6C, 6E-G**) at a pixel size of 0.415 µm X 0.415 µm or a 10X Plan Apochromat objective (NA: 0.45; WD: 2.0 mm; (**Figure 6D**)) at a pixel size of 1.661 µm X 1.661 µm. A fixed pinhole size at 1 Airy unit based on the 551/632 channel was used for multi-channel imaging. Additional images (**Figure 4J**) were acquired using an Olympus FV3000RS confocal microscope equipped with four fluorescence spectral detectors. Excitation lasers used were: 405 nm for DAPI, 488nm for Alexa Fluor 488, 543 nm for Alexa Fluor 568 and Cy3, and 633 nm for Alexa Fluor 647. Emission filters used were: 406/545 for DAPI and Alexa Fluor 488, 551/632 for Alexa Fluor 568 and Cy3, and 637/756 for Alexa Fluor 647. Original images were obtained using a 10X Super Apochromat objective (NA: 0.4; WD: 3.1 mm) at a pixel size of 2.49 µm x 2.49 µm. A fixed pinhole size at 1 Airy unit based on the 551/632 channel was used for multi-channel imaging.

### Confocal Image Processing and Cellular Quantification

Confocal images were processed and analyzed using FIJI (v2.16.0). To quantify cell populations within the P14 cortex, multi-channel images were first separated into individual monochromatic channels (DAPI, NeuN, Olig2, and Iba1). To ensure unbiased quantification, all image processing and cell counting was performed by a researcher blinded to the genotype of the samples. For each channel, a global threshold was manually determined to maximize the signal-to-noise ratio and binarize the image. To resolve overlapping nuclei, a distance-transform-based watershed segmentation algorithm was applied to the binary masks. Automated cell counting was performed using the ‘Analyze Particles’ function. To ensure the exclusion of non-specific labeling and physical artifacts, a size-exclusion filter was applied. Total cell counts for each marker were normalized to the total DAPI+ population to calculate the relative percentage within the cortex. All processing parameters were kept consistent between WT and *Chd8^V986*/+^*mice to ensure unbiased quantification.

### Manual Nuclei Annotation

Recent deep learning-based nuclear segmentation algorithms show great promise for automated segmentation, but require large numbers of accurate manually labeled nuclei as either training data or validation of accuracy. We developed a nuclei annotation tool as a stand-alone app called Segmentor^114^ and a web-based tool that enables crowdsourcing of nuclei annotation work for citizen science called ninjato^115^. We used these tools to annotate the full 3D shape of 10,829 nuclei in lightsheet images of the mouse brain at both P4 and P14 (**Table S12**). These full 3D nuclei annotations were then used to train our NIS model described in the next section.

### Nuclei Instance Segmentation (NIS)

Our NIS model predicts the probability that a voxel would be a nucleus instance along with 3D spatial gradients based on a 2D Unet. Cellpose as a general cell instance segmentation method has a 3D approach in its first version^42^. However, this 3D method is inefficient since it repeats the whole-brain computation on xy, xz, and yz planes three times, tripling the computing time. Our method design is inspired by Cellpose^42^. We first reuse the 2D method of Cellpose, training a Unet supervised by spatial gradients in 2D planes that have a higher resolution, i.e., 0.75 μm per voxel (**Figure S6A**). We then propose a median filter pyramid (MFP), rebuilding the spatial gradient in the third dimension (4 μm per voxel) during the inference, to avoid tripled computing time required by 3D Cellpose 1.0^42^. MFP as the 3D recon block in **Figure S6B** has a dynamic estimation taking the L2-norm of 2D gradient from the previous and the next 2D slices. We initialize the third dimension gradient as the next slice gradient norm minus the norm in the previous slice. For voxels with similar norms inside the nucleus, we iteratively update the gradient norm map using a 3×3 median kernel, and average the difference between the filtered norm from every iteration as the third gradient, where nuclei with variable sizes can be gradually covered by repeating the iteration 7 times. Then, we threshold the probability map and stack 2D maps along the third dimension as 3D maps, only considering probabilities larger than zero. To accomplish NIS reconstruction (denoted by ‘recon block’ in **Figure S6B**) given the thresholded 3D maps, we reuse the dynamic system of Cellpose 1.0^42^ by starting at every foreground voxel and following the spatial derivatives specified by the 3D gradient. Each time a step was taken towards the direction of the gradient at the nearest grid location within the foreground. Following convergence, voxels can easily cluster according to the location they end up at. For robustness, the clusters are extended 3 voxels along regions of high-density foreground convergence to prevent the exact center from being missed by Unet in some cases.

Due to limited computational resources, it is a common practice to split the full image into a set of non-overlapping chunks and apply NIS separately within each chunk. For the NIS coordinates located at the cropping boundary of the 3D maps, we utilized our NIS stitching method^116^, by building a graph for cropped NIS masks and train graph neural networks to stitch, denoted by Graph Construct and GCN (Graph Convolution Networks) blocks in (**Figure S6B**). The coordinates located at the cropping boundary of each cluster are first collected as a node of the graph, where each node is assigned a vector of 227 elements as the node feature. The first 200 elements are the resampled image intensities within the NIS mask, and the last 27 elements are the counts of 27 (3x3x3) possible directions of 3D spatial gradient. Edges of the graph are connected to the top 2 closest nodes and are assigned with the spatial distance between centers, height and width ratios, and the average spatial gradient. GCN takes this graph as input and classifies if the edges connect the same nuclei as the output. The training of GCN is supervised learning via cross-entropy loss between the output and the annotation generated from the 3D segmentation mask. Every two neighboring slices of the 3D mask were used to generate the annotation. GCN was trained for 300 epochs with SGD with a learning rate of 0.001, a momentum of 0.9, a batch size of 64 pairs of neighboring mask slices, and a weight decay of 0.00001. The learning rate started from 0.001 and was linearly annealed with a weight of 0.5 every 30 epochs. For the NIS coordinates located at the overlapping area of two neighboring tiles, we implemented two approaches to alleviate double counting: (1) a coarse-to-fine wobbly stitching and (2) undoubling the segmentation by removing overlapping NIS. The wobbly stitching starts from a phase correlation correction (PCC) between every pair of 3D tiles to estimate a coarse stitching transformation for the entire 3D tile, and then applies the average transformation from all neighbors to each 3D tile. The coarse stitching has to load whole-brain LSFM images, resulting in 2 hours of computational time. Next, the Coherent Point Drifting (CPD) registration method^117^ is applied as refined stitching every 25 slices between a coarsely stitched tile and all neighboring refined tiles using nuclei in these regions as reference points. In addition to the two-stage automatic stitching, manual stitching utilizing Imaris Stitcher was used as a post-hoc step for failed cases. Briefly, 3D image tiles were aligned and fused using Imaris Stitcher (v10.2.0). Overlapping regions were manually registered using the software’s “drag-and-drop” tool. The resulting stitched images and stitching parameters were exported as .tif and .xml files for subsequent analysis.

The training of Unet is implemented based on codes of Cellpose 2.0^118^ upon 3,227 and 6,032 Topro3 nuclei from 5 P4 and 4 P14 brains, respectively, which were manually annotated using Segmentor^114^ and Ninjato^115^. In addition, 755 and 815 Topro3 nuclei from 1 P4 and 2 P14 brains, respectively, were used for the segmentation accuracy evaluation. Precision, recall, and F1 score are metrics. The definition of True Positive is 3D IoU (Interaction over Union) ratio between the predicted mask and the ground truth mask over 0.3. Unet was trained for 300 epochs with SGD with a learning rate of 0.001, a momentum of 0.9, a batch size of 64 128×128 images, and a weight decay of 0.00001. The learning rate started from 0.001 and was linearly annealed with a weight of 0.5 every 30 epochs. The NIS visualization at the whole-brain scale is implemented by the density map with a lower resolution. The downsampled density map reduces the memory requirement while preserving the precise counts of NIS for downstream analysis by assigning the number of nuclei centers within the local cubic area (25x25x25 μm^3^) to the value of each voxel in the density map.

### Colocalization

The co-localization on multiple channels, i.e., Brn2 and Ctip2, and NeuN and Sox9 of P4 and P14 brains, respectively, was achieved using NIS-guided ResNet50^119^. Considering the unbalanced SNRs between different channels, the global location of a cell is an important prior knowledge for the manual annotation. We implemented a Matlab annotation tool to display 100 to 1,000 local patches of NIS results aside with a whole-brain location indicator as shown in **Figure S6F** top part. Local patches can be randomly picked or explicitly selected from interesting regions with a brain mask, which is created for each individual brain where errors were found during the quality check. As a result, we conducted 18,235 and 6,228 patch images of Brn2/Ctip2 and Neun/Sox9 cells from 32 P4 brains (M/F=16/16, WT/HET=16/16) and 17 P14 brains (M/F=7/10, WT/HET=8/9), respectively, bringing total numbers of manually annotated cell types as listed in **Table S12**. Then, authors FK and IC manually determined the patch label based on the information of multichannel intensity and whole-brain location. For either training or inference as shown in **Figure S6F** bottom, the patch image of each channel is first normalized to [0,1], and then stacked as a multichannel LSFM image, cropped by the NIS bounding box, and resized to 30×30. During the training, there are three image augmentations randomly (50%) applied to the image for training data variability. Additionally, (i) the spatial location (x, y, z) is embedded and concatenated to the latent feature from the patch, and (ii) the imbalanced sampling is applied according to the cell count per channel, which is uneven across the whole brain. The training of ResNet50 has 500 epochs and 1e-4 learning rate using Adam optimizer^119,120^. For inference, the image augmentations were skipped. For Brn2/Ctip2 multichannel LSFM images, training cells were selected in subregions of the isocortex where upper and lower layer neurons mainly existed.

#### Heterotopia Detection in *Chd8* Exon 3 Mutation Model (*Chd8^ΔEx3/+^*; Suetterlin et al., 2018)

All procedures were done in accordance with the Animals (Scientific Procedures) 1986 Act with ethical approval granted by the King’s College London Animal Welfare and Ethical Review Body and the UK Home Office. Mice were housed under a 12-hour light cycle, with ad libitum access to food and water. Constitutive *Chd8^ΔEx3/+^* mice were generated as previously described^48^ and maintained by crossing *Chd8^ΔEx3/+^* and C57BL/6J mice. At 7 postnatal weeks, animals were deeply anaesthetised with pentobarbital and transcardially perfused with 4% PFA/PBS. Brains were extracted and post-fixed overnight in 4% PFA at 4°C. Brains were sectioned using a Campden instruments 5100mz vibratome at 70 µm/section, for a total of 28-35 sections starting from the frontal part of the brain. Coronal sections were collected in PBS + 0.02% sodium azide and stored at 4°C before immunofluorescence staining. Sections were blocked in PBS supplemented with 0.25% TritonX-100 and 10% normal goat serum for 2 hours at room temperature, then incubated overnight at 4°C in primary antibody diluted in blocking solution. Brain slices underwent three 10-minute washes in blocking solution, followed by a 1-hour incubation in secondary antibody solution at room temperature. Sections were washed three times for 10 minutes in PBS and then counterstained with DAPI (1:5000 in PBS) for 10 minutes before another 10-minute wash with PBS. Sections were mounted onto standard microscope slides (Academy, 7107, 1.0-1.2mm thick) with Fluoroshield Anti-Fade mounting medium (Abcam, ab104135) and stored at 4°C. The following antibodies were used: mouse anti-NeuN (Sigma-Aldrich, MAB377, 1:250) and goat anti-mouse IgG Alexa Fluor 568 (Thermo Fisher Scientific, A11031, 1:1000). From the total pool of collected tissue, a representative subset of 7-14 coronal sections per animal was selected for further analysis. Images were acquired using a Zeiss Axioscan slide scanner. Excitation LEDs used were: 385 nm for DAPI and 557 nm for Alexa Fluor 568. Emission filters used were: 450/40 for DAPI and 605/70 for Alexa Fluor 568. Beam splitters used were 420 and 570. Original images were obtained using a 10X Plan Apochromat objective (NA: 0.45; WD: 2.0 mm) at a pixel size of 0.455 µm X 0.455 µm. Heterotopias were detected manually in QuPath. Differences in proportions of animals with heterotopias between groups were evaluated using a two-tailed Fisher’s exact test.

#### Heterotopia Detection in *Chd8* Exon 5 Mutation Model (*Chd8^5bpdel/+^*; Gompers et al., 2017)

Generation and initial characterization of the Cas9-mediated 5 bp frameshift deletion in exon 5 of *Chd8* mice (here referred to as *Chd8^5bpdel/+)^* were previously described^19^. A total of 32 mice were included in the present study (16 *Chd8^5bpdel/+^* and 16 *wild-type* littermates; balanced by sex). Animals were analyzed at postnatal day 42. All experimental procedures were approved by the Institutional Animal Care and Use Committee (IACUC) at the University of California, Davis and were conducted in accordance with institutional guidelines. Mice were housed under standard conditions in a temperature-controlled vivarium on a 12 h light/dark cycle with ad libitum access to food and water. All efforts were made to minimize animal distress and reduce the number of animals used. Mice were deeply anesthetized with isoflurane and transcardially perfused with 0.9% saline followed by 4% paraformaldehyde (PFA) in 0.1 M phosphate buffer (PB) using a peristaltic pump (Minipuls 3, Gilson; ∼20 rpm), with perfusate volumes scaled to body weight (∼1 mL per gram). Brains were removed, post-fixed in 4% PFA at 4 °C for 24 h, and cryoprotected sequentially in 10% glycerol/2% DMSO (24 h) and 20% glycerol/2% DMSO (72 h) in 0.1 M PB (pH 7.4). Prior to freezing, each brain was blocked at −2.00 mm posterior to the interaural line using an adult mouse brain matrix (Zivic Instruments), removing the cerebellum and caudal brainstem; the resulting block therefore spanned the full rostro-caudal extent of the cerebrum. Blocks were flash-frozen on dry ice and stored at -70 °C until sectioning. Coronal sections were cut at 60 µm on a freezing sliding microtome (Microm HM 440E, Microm International) and collected as four interleaved serial sets, such that adjacent sections within any one set were spaced 240 µm apart (4 × 60 µm). One complete series (every fourth section) was post-fixed in 10% formaldehyde in 0.1 M PB for 1 week at 4 °C and processed for Nissl staining; the remaining three series were stored in cryoprotectant at -70 °C. Because collection spanned the entire blocked cerebrum, the Nissl series sampled the brain throughout its rostro-caudal extent (excluding the cerebellum) rather than being restricted to any single region. For staining, sections were rinsed in 0.1 M PB, mounted onto gelatin-coated slides, and dried overnight at 37 °C. Tissue was defatted in chloroform/ethanol (1:1) for 2 × 1 h, rehydrated through graded ethanol (100%, 95%, 70%, 50%) to distilled water, stained in 0.25% thionin (Fisher Scientific, T-409) for 40 s, differentiated in 95% ethanol containing glacial acetic acid, dehydrated through graded alcohols, cleared in xylene, and coverslipped with DPX mounting medium (BDH Laboratories). The

complete Nissl-stained series from each animal was systematically screened for molecular layer heterotopias (MLHs) along the full rostro-caudal extent of the cerebrum (cerebellum excluded). Screening was performed under blinded conditions by two independent investigators to minimize observer bias. Sections were examined by brightfield (transmitted-light) microscopy on an Olympus SZX16 stereomicroscope at low magnification (10×; LED illumination), and images were acquired with an Olympus DP23 color camera (6.4-megapixel CMOS image sensor) using Olympus cellSens Entry v3.2 software.

### Heterotopia Detection in C57BL/6J x DBA/2J (B6-DBA) F1 hybrids

C57BL/6J x DBA/2J (B6-DBA) F1 mutant and littermate control mice were generated by breeding male *Chd8^V986*/+^*mutant mice in C57BL/6J background with wild-type females in DBA/2J background (The Jackson Laboratory; Strain #:000671; RRID:IMSR_JAX:000671). Pups were perfused at the age of one month (P30) and brains were dissected and postfixed as described above. Serial coronal sections at 50µm were collected from the rostral tip of the cortex to or beyond the level of Bregma -1.0. All brains sections were subjected to immunofluorescence staining against NEUN and DAPI staining to detect heterotopias. Guinea pig anti-NEUN (Millipore, ABN90P; 1:1,000) were used as the primary antibody and Alexa Fluor 568 Donkey Anti-Guinea Pig IgG (Jackson ImmunoResearch, 706-575-148; 1:1,000; RRID: AB_3095468) were used as the secondary antibody. Presence of heterotopias were visually evaluated using a Zeiss Axioskop fluorescence microscope equipped with a 20X LD Achroplan objective (NA: 0.40; WD: LD; Korr: 0 – 1.5 mm) and an EXFP X-Cite 120 PC fluorescence illumination system. Observed heterotopias were confirmed by confocal imaging using a Zeiss LSM 710 laser scanning confocal microscope equipped with three fluorescence spectral detectors. Excitation lasers used were: 405 nm for DAPI, and 543 nm for Alexa Fluor 568. Emission filters used were: 406/545 for DAPI and 551/632 for Alexa Fluor 568. Beam splitters used were MBS 488/543/633 and MBS 405. Z-stack images (z = 10µm; step = 1 µm) were obtained using a 63X Plan Apochromat objective (NA: 1.4/Oil; WD: 0.17 mm) at a pixel size of 0.264 µm X 0.264 µm. A fixed pinhole size at 1 Airy unit based on the 551/632 channel was used for multi-channel imaging. Resulting images following maximum intensity projection were presented in Figure 6H.

### snRNA-seq in *Chd8^5bpdel/+^* embryos

snRNA-seq data was obtained from the *Chd8^5bpdel/+^* mouse line^19^. After timed breeding, embryos were removed 14.5 days post conception, and the forebrain was dissected out and flash frozen for snRNA-seq. 2 litters with balanced sex and genotype composition were used, with a final n = 4 per genotype. The 10X Genomics nuclei isolation kit, with protocol as described, was used to isolate nuclei. To limit batch effects WT and HET mice were combined into pools, each containing one male and one female of each genotype, which was deconvoluted in downstream analysis. Nuclei suspensions were counted and adjusted for max 10X library targeting, then used as input for the 10X Genomics Multiome Assay. Libraries were generated according to the v3 protocol by the UC Davis DNA Technologies Core. 10X libraries were sequenced with the Element Biosciences AVITI at 80,000 reads per nucleus. De-multiplexed sequencing data were obtained from the UC Davis DNA Technologies Core as FASTQ files. FASTQ files were passed through the Cell Ranger *arc* pipeline for alignment to the mouse mm10 genome, filtering, and creation of feature-barcode matrices. Ambient RNA removal was performed using SoupX, and Seurat objects

were generated using Seurat. These Seurat objects were subjected to initial clustering, and each resulting cluster was filtered based on cluster-specific thresholds for UMI count, feature count, and percentage of mitochondrial reads, which were determined using standard deviation criteria. Following initial QC, doublets were further filtered out with the scDblFinder package. Sex and genotype were then assigned to each cell based on expression of X and Y-linked genes. All sequencing batches were normalized with SCTransform and integrated with RPCAIntegration. Standard nearest neighbors and clustering methods were followed, and dorsal pallial radial glia clusters were identified via expression of Pax6 and other canonical markers. We obtained 795 high quality pallial radial glia cells for differential expression analysis. Gene counts from radial glial cells were aggregated within biological replicates to generate pseudobulk expression profiles for differential expression analysis. Differential expression was performed using DESeq2 with sex as a covariate, which employs negative binomial generalized linear models to estimate gene-level effect sizes and statistical significance between conditions. To reduce variance associated with lowly expressed genes, only genes with at least 10 counts in at least half of the samples were retained for analysis.

Differential gene expression and pathway enrichment analysis (*Chd8^A408fsX/+^*; Yim et al., 2024)

We obtained single-cell RNA-seq (scRNA-seq) from *Chd8^A408fsX/+^*and WT developing mouse cortex from previously published data (GEO accession GSE273271)^26^. The raw gene expression matrices were reanalyzed following the author’s publicly available GitHub repository (https://github.com/NoonanLab/Yim_et_al_Chd8), with minor modifications. Briefly, QCed data was processed using Seurat V5^121^. Data split by samples (batch, sex and genotype (WT vs *Chd8^A408fsX/+^*) were merged using the JointLayer() function. Clustering was performed by FindClusters() at resolution 1.2 to identify finer cell type classification. For differential expression, we applied a pseudobulk strategy. Counts were aggregated per sample and per cell type using the aggregateData() function from the muscat package (v1.22.0)^122^. Differential expression between wild-type and *Chd8^A408fsX/+^*at each time point were tested using limma-voom^123,124^ while correcting batch and sex. Gene with Bonferroni-adjusted p-values < 0.1 were considered significant. Gene set enrichment analyses were performed using the fgsea package (v1.34.2)^125^ only on radial glia cell type (cluster = 3) at E12.5. Genes were ranked by -log(Bonferroni-adjusted p-values) x signed(logFC) to incorporate both significance and direction of regulation.

### *Chd8^+/-^* differential gene expression in radial glial endfeet analysis

We obtained the background list from all genes with reads detected in the E12.5 scRNAseq *Chd8^A408fsX/+^* DEGs as described in Figure 7K or E14.5 snRNAseq *Chd8^5bpdel/+^* DEGs as described in Figure S11 and bulk RNA-seq of radial glial endfeet and cell bodies^54^. Each gene was designated as endfoot localized and/or *Chd8* differentially expressed using WT vs *Chd8^A408fsX/+^*cluster 3 cells at E12.5 in Figure 7K and radial glia in WT vs *Chd8^5bpdel/+^*at E14.5 in Figure S11. Statistical significance of the overlap between DEGs and endfoot localized genes was tested using a two-tailed Fisher’s exact test.

### Statistics for MRI Volume

All 3D volume analyses and regional isocortex analyses were conducted using custom R scripts. P values for Total Brain Volume and Isocortex combined regions were estimated from a multiple linear regression controlling for pair (equivalent to a paired t-test): Volume ∼ Genotype + Pair. Statistical significance was reached at p<0.05. Regional summary statistics were computed using the 17-region isocortical parcellation (Allen reference atlas; depth = 6). Following registration and segmentation, measurements were aggregated according to the Allen CCFv3 ontology tree, first collapsing by cortical layer and then by higher-level cortical areas to obtain the 17 regions corresponding to the immediate children of the isocortex structure. For the 17 regional cortical regions, false discovery rate (FDR) adjusted p values (Benjamini-Hochberg; FDR < 0.1) were extracted from the same multiple linear regression controlling for pair.

### Statistics for Cell Counts, Colocalization, and Density

All cell counts analyses and regional isocortex analyses were conducted using custom R scripts. P values for Isocortex combined regions were estimated from a multiple linear regression controlling for pair and manually annotated tissue loss: Counts ∼ Genotype + Pair + Tissue loss. Statistical significance was reached at p<0.05. For the 17 regional cortical regions, FDR adjusted p values (Benjamini-Hochberg; FDR < 0.1) were extracted from a multiple linear regression controlling for pair and tissue losses: Counts ∼ Genotype + Pair + Tissue loss. The 17-region isocortical parcellation was generated from the Allen reference atlas as described above.

### Visualizations of Heterotopia Detection

To ensure all heterotopias were detected at cellular resolution, we partitioned our large light-sheet datasets into 200um thick subsections. These subvolumes were imported into ITK-SNAP (v4.4.0) for systematic visual inspection of the pial surface. This approach allowed the visualization of the cortical plate at full resolution, preventing missed malformations due to computational down-sampling. Heterotopias were identified and manually segmented using the polygon tool in ITK-SNAP to generate 3D binary masks. These masks were used for subsequent volumetric quantification and spatial visualizations.

### Brain Renderings

For visualizing volumetric data, VTK mesh surfaces of the Allen Reference Atlas and individual MRI scans were generated using custom Python code. Mesh generation relied on isosurface extraction via the Marching Cubes algorithm implemented in *scikit-image* package. The resulting surfaces were reconstructed and smoothed in PyVista. The same workflow was used to generate 3D surface meshes of heterotopia for morphological inspection.

For region-based analyses, cortical contours were derived from Allen atlas label maps. During mesh generation, brain-region labels from either the original atlas or the registered subject-specific label image were propagated to mesh vertices using a nearest-neighbor assignment. ParaView’s Python interpreter (PvPython) was then used to generate contours that delineate boundaries between adjacent regions. Specifically, evenly spaced boundary lines were computed between regions and then extracted as iso-surfaces, which were assembled into a final contour surface mesh and exported as a VTK file.

Regional summary statistics were computed using the 17-region isocortical parcellation (Allen reference atlas; depth = 6). The 17-region isocortical parcellation was generated from the Allen reference atlas as described above. Regional metrics were then mapped onto the contoured surface mesh for visualization.

### Point clouds

VTK surface renderings were also used to visualize cell-density distributions after nuclei segmentation. To improve memory efficiency, cell coordinates were rescaled and aligned to the target voxel resolution of the atlas or density map. Cells were then randomly subsampled at a user-defined proportion to reduce overplotting. A binary mask derived from mesh vertex labels was applied to retain only cells within the isocortex. Surface transparency was adjusted in the PyVista plotter to achieve a “glass brain” effect. Example scripts demonstrated manual camera positioning and export of high-resolution renderings.

### Flatmaps

A flattened isocortex representation was used to visualize shifts in cell counts and densities between heterozygous and control groups This visualization was adapted from the flatmap approach used in NuMorph, which uses a precomputed mapping between 3D cortical surface coordinates and a 2D projection^31,126^. The 2D flatmap included the same 17-region isocortical contour configuration.

## Author Contributions

Study Design: FAK, IC, ZW, GW, JLS P4 MRI and lightsheet microscopy: FAK

E18.5 and P14 MRI and lightsheet microscopy: IC

Development and application of nuclei instance segmentation and colocalization: ZW Statistical analyses: FAK, IC

3D visualizations: MYY

Immunohistochemical characterization of *Chd8*^V986*/+^ in slices: LX, IC, FAK, BTB Dual view deconvolution on E18.5 images: TW

Image registration: RMV, MD Manual image annotations: KL scRNA-seq analysis: NM, NS

Animal breeding: CMM, TF, IC, FAK, ESM

Tissue clearing protocol optimization: IC, FAK, OK, MRG MRI acquisition: TWW, QH, YIS

Radial glial endfoot localized genes analysis: BRD, DLS Manual segmentation tool development: DB, HY

Manual nuclei segmentations: LTD, VD, CE-T, KE, MM, MY, KH, MB, MK Light sheet imaging optimization: PA

Processing, imaging, and snRNA-seq of *Chd8^5bpdel/+^* mice: CPC, NS, JB, DGA Processing and imaging of *Chd8^ΔEx3/+^* mice: PPT, LP-S, LCA

Development of *Chd8*^V986*/+^ mouse line: MJZ Development of *Chd8^5bpdel/+^* mouse line: ASN

Development of *Chd8^ΔEx3/+^* mouse line: MAB

Joint supervision of work: JLS, GW, MJZ, ASN Acquisition of funding: JLS, GW, MJZ, ASN

## Data Availability

The raw MRI and multi-channel light sheet images supporting the findings of this study have been deposited in the Brain Observatory Storage Service and Database (BossDB) and are publicly available at https://bossdb.org/project/curtin2026.

## Code Availability

NIS, stitching, and co-localization codes are available in https://github.com/Chrisa142857/Lightsheet_microscopy_image_3D_nuclei_instance_segmentation. All statistical analysis and visualization codes are available at https://bitbucket.org/FelixKyere/chd8_project_image_analysis/src/master/.

## Supplementary Materials

Due to file sizes, Videos S1 to S5 are uploaded on YouTube at the specified links below: Materials and Methods

Supplementary Text Figures S1 to S11 Tables S1 to S12

Video S1: https://youtu.be/GTXCTUQTUyA Video S2: https://youtu.be/DKY3SyZmLzk Video S3: https://youtu.be/62xBlLmnIWg Video S4: https://youtu.be/QMHHBEd0b80 Video S5: https://youtu.be/FLZlYHxe-gg

## Supporting information

Raw values of MRI, NIS, Tissue loss and Density for WT and Chd8V986*/+ samples at P4.

Raw values of MRI, NIS, Tissue loss and Density for WT and Chd8V986*/+ samples at P14.

Raw values of colocalization Density across cortical regions for all samples

Summary of E18.5, P4 and P14 samples used for each analyses across multiple imaging modalities.

Summary statistics of colocalization with TO-PRO-3 across cortical regions between WT and Chd8V986*/+ million counts/mm3.

Raw values of colocalization with TO-PRO-3 across cortical regions for all samples

Summary statistics of colocalization with TO-PRO-3 across cortical regions between WT and Chd8V986*/+

Summary statistics of density analyses across cortical regions between WT and Chd8V986*/+

Raw values of tissue shrinkage and summary statistics for WT and Chd8V986*/+ mice

Number of training data for NIS (sheet 1) and colocalization (sheet 2) organized by sex, genotype and time point.

Summary statistics of volumetric analyses across cortical regions between WT and Chd8V986*/+

Summary statistics of nuclei segmentations across cortical regions between WT and Chd8V986*/+

Representative Images of all markers of WT and Chd8V986*/+ brains at P4.

Representative Images of all markers of WT and Chd8V986*/+ brains at P14.

3D Volumetric Reconstruction and 2D Cross-Sectional Views of Molecular Layer Heterotopia in E18.5 Chd8V986*/+ brain.

3D Volumetric Reconstruction and 2D Cross-Sectional Views of Molecular Layer Heterotopia in P4 Chd8V986*/+ brain.

3D Volumetric Reconstruction and 2D Cross-Sectional Views of Molecular Layer Heterotopia in P14 Chd8V986*/+ brain.

## Acknowledgements

The Microscopy Services Laboratory, Department of Pathology and Laboratory Medicine, is supported in part by P30 CA016086 Cancer Center Core Support Grant to the UNC Lineberger Comprehensive Cancer Center. The Neuroscience Microscopy Core is supported by P30 NS045892. The Center for Animal MRI is supported in part by S10 OD026796, S10 MH124745, P60 AA011605, and P50 HD103573. Research reported in this publication was also supported in part by the North Carolina Biotech Center Institutional Support Grant 2016-IDG-1016. The authors gratefully acknowledge BossDB.org for providing data storage and access services that support the sharing of the datasets associated with this publication (R24MH114785, Hider et. al 2022 - https://doi.org/10.3389/fninf.2022.828787). We thank Tim Bell, Matt Blanchard, and Fernando Pardo Manuel de Villena for performing MiniMuga genotyping and analysis. This work was supported in part by the Regenerative Medicine Development Organization (ReMDO) Test Bed, which provides infrastructure and translational development resources to accelerate regenerative medicine innovation. The ReMDO Test Bed is supported in part by Wake Forest Institute for Regenerative Medicine (WFIRM), the Health & Services Administration (HRSA), the National Science Foundation (NSF) Regional Innovation Engines award, and the Piedmont Triad Regenerative Medicine Engine.

