## Supplemental Figures for "*Chd8* haploinsufficiency leads to molecular layer heterotopias and age-dependent cortical expansion"

Affiliations:
[1]UNC Neuroscience Center, University of North Carolina, Chapel Hill, NC, 27599, USA.

[2]Department of Genetics, University of North Carolina, Chapel Hill, NC, 27599, USA.

[3]Department of Computer Science, University of North Carolina, Chapel Hill, NC, 27599, USA.

[4]UC Davis Center for Neuroscience, Davis, CA 95618, USA.

[5]Department of Psychiatry and Behavioral Sciences, University of California Davis, Sacramento, CA 95817, USA.

[6]The M.I.N.D. Institute, UC Davis, Sacramento, CA, USA.

[7]Centre for Developmental Neurobiology, Institute of Psychiatry, Psychology and Neuroscience, King's College London, London, United Kingdom; MRC Centre for Neurodevelopmental Disorders, King's College London, United Kingdom.

[8]Department of Psychiatry, University of North Carolina, Chapel Hill, NC, 27599, USA.

[9]Center for Animal Magnetic Resonance Imaging, University of North Carolina, Chapel Hill, NC, 27599, USA.

[10]Biomedical Research Imaging Center, The University of North Carolina at Chapel Hill; Department of Neurology, The University of North Carolina at Chapel Hill, Chapel Hill, NC, 27599, USA.

[11]Department of Cell Biology, Duke University School of Medicine, Durham, NC, 27710, USA.

[12]Departments of Molecular Genetics and Microbiology and Neurobiology, Duke University School of Medicine, Durham, NC, 27710, USA.

[13]Duke Institute for Brain Sciences and Duke Regeneration Center, Duke University School of Medicine, Durham, NC, 27710, USA.

[14]RENCI, University of North Carolina at Chapel Hill, 100 Europa Drive, Suite 540, Chapel Hill, NC 27517 USA.

[15]Microscopy Services Laboratory, Department of Pathology and Laboratory Medicine, University of North Carolina at Chapel Hill, 27599, USA.

[16]Clinical and Biomedical Sciences, University of Exeter Medical School, Hatherly Laboratories, Exeter EX4 4PS, U.K.

[17]Carolina Institute for Developmental Disabilities, University of North Carolina, Chapel Hill, NC, 27599, USA.

[18]Department of Cell Biology and Physiology, University of North Carolina, Chapel Hill, NC, 27599, USA.

[19]Lead contact

† Contributed equally
†† Contributed equally
‡ Jointly supervised

a

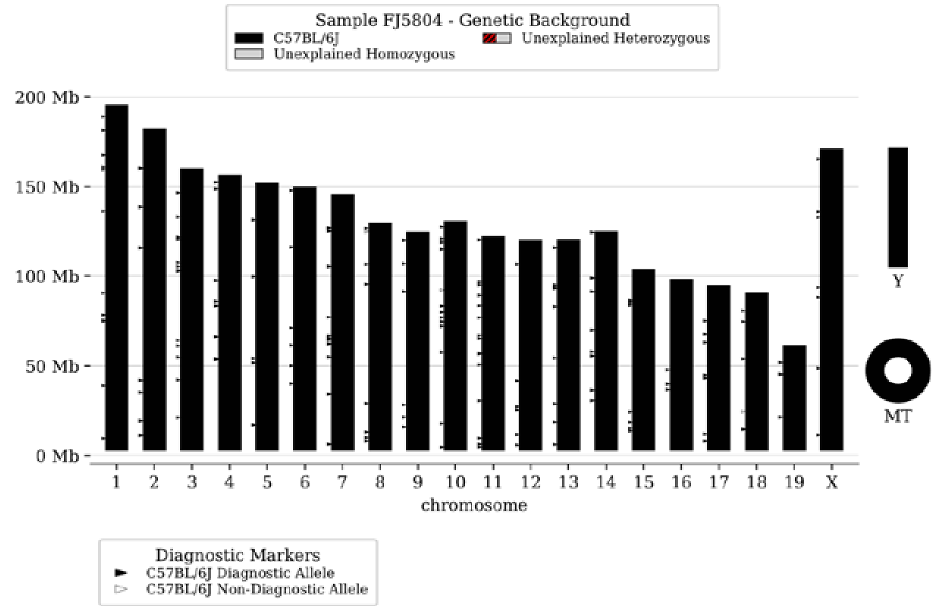


b

| | |
|---|---|
| **BlastR** | - |
| **bpA** | - |
| **Cas9** | - |
| **chlor** | - |
| **cHS4** | - |
| **Cre** | - |
| **DTA** | - |
| **Flp** | - |
| **g_FP** | - |
| **hCMV_a** | - |
| **hCMV_b** | - |
| **hTK_pr** | - |
| **iCre** | - |
| **IRES** | - |
| **Luc** | - |
| **r_FP** | - |
| **rtTA** | - |
| **SV40** | - |
| **tTA** | - |

**Figure S1: Genetic Validation of the C57BL/6J Mouse Strain.**

To confirm the genetic background of the experimental mouse line, a MiniMUGA genotyping panel was performed on eight mice across four littermate pairs (n WT = 4 and n *Chd8*$^{V986*/+}$ = 4). This cohort included balanced sex distribution (4 males, 4 females) and represented mice both with (n = 2) and without (n = 6) heterotopias.

a: Representative karyoplot of the genetic background analysis. All 19 autosomes and the X chromosome were identified as 100% homozygous (or hemizygous) for all C57BL/6J mice. Analysis of diagnostic alleles confirmed the sample is consistent with the C57BL/6J strain (165/168 markers, 98.2%).

b: Representative summary of genetic construct analysis. All samples were confirmed to be 100.0% inbred status for both males (XY) and females (XX) with "Excellent" genotyping quality (7 N calls). All samples tested negative for all 19 common genetic constructs, including Cre, Flp, and Cas9.

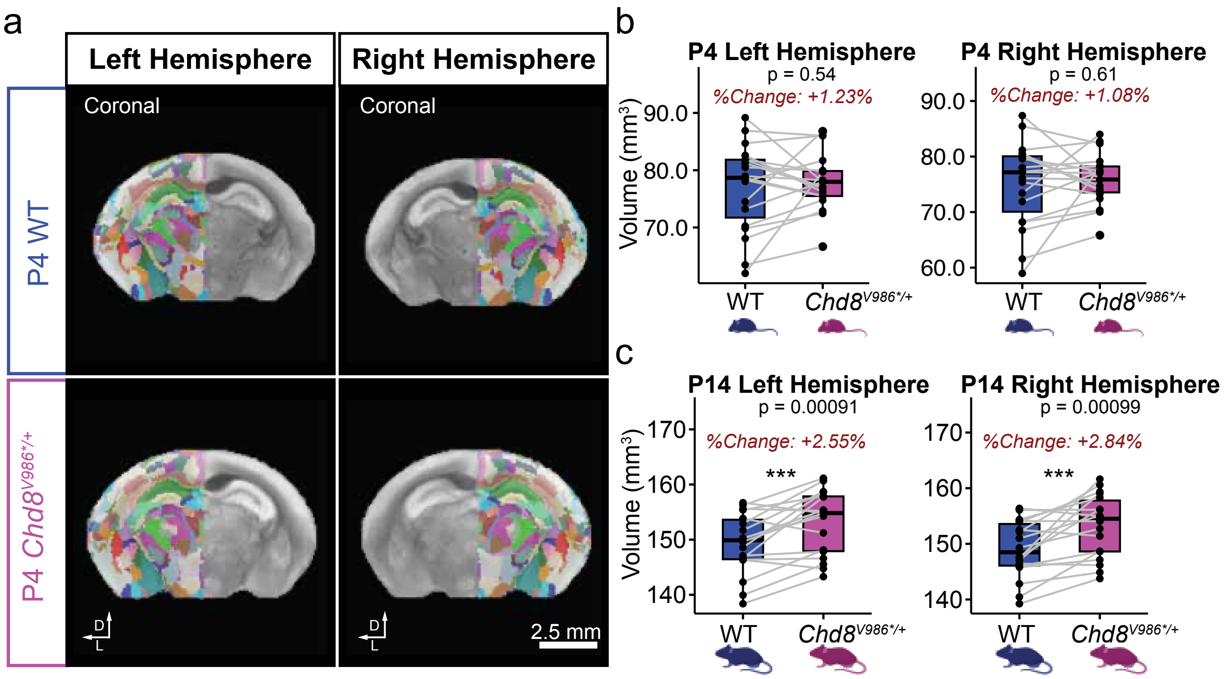


**Figure S2: Brain Volume Laterality Effects.**

a: Representative images for WT (top row) and *Chd8*$^{V986*/+}$ (bottom row) showing the left and right MRI hemispheres at P4. Overlap on either left or right hemisphere shows atlas registration.

b: Volume measurement of left hemisphere (left box plot) and right hemisphere (right box plot) at P4. P-value was evaluated using a multiple linear regression controlling for pair ($n_{pairs}$ = 19)

c: Volume measurement of left hemisphere (left box plot) and right hemisphere (right box plot) at P14. P-value was evaluated using a multiple linear regression controlling for pair ($n_{pairs}$ = 18; *** p < 0.001).

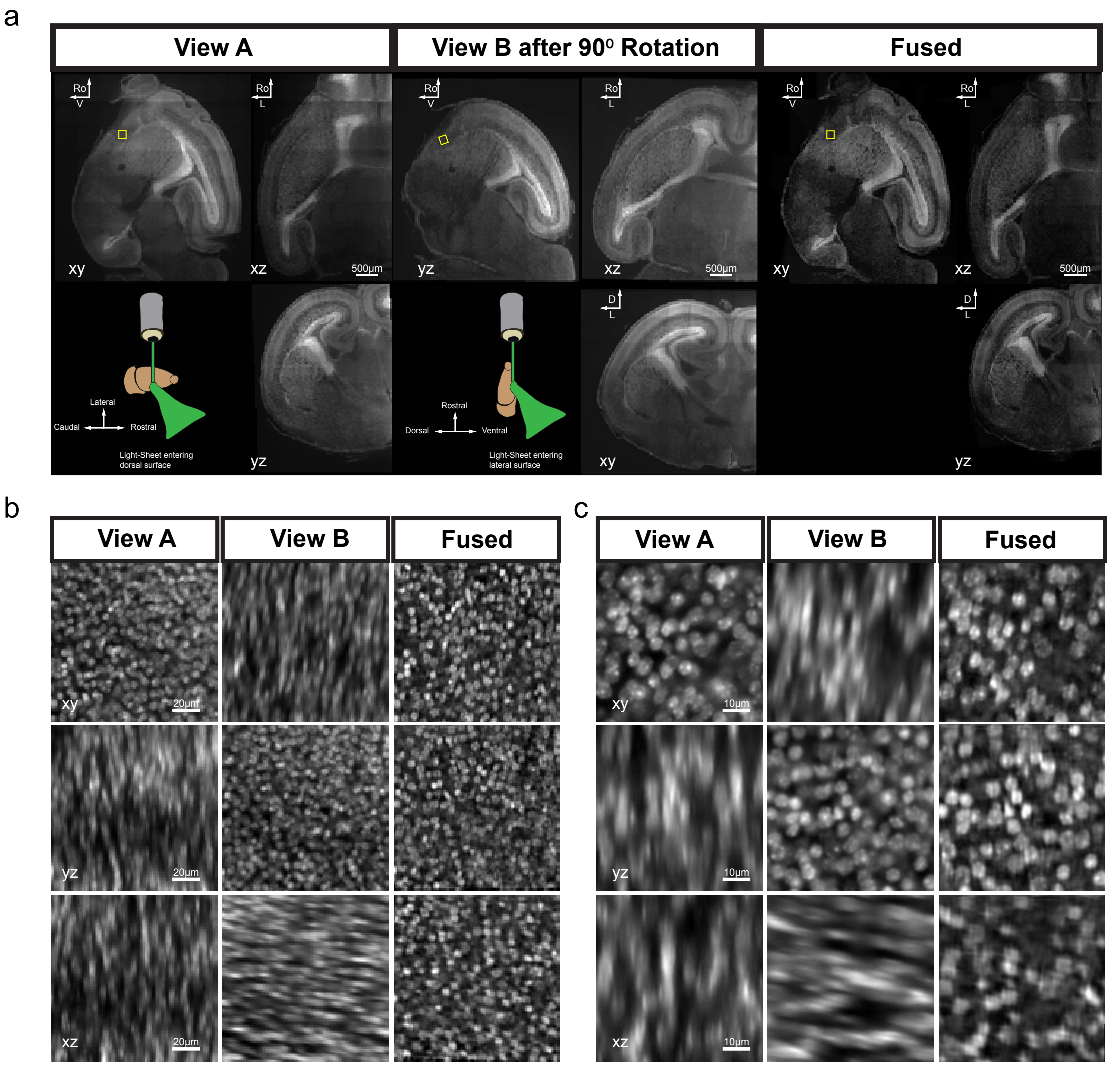


**Figure S3: Multi-view Cellular-scale Imaging with Sample Rotation and Anisotropy-resolved Fusion.**

a: Large-scale tissue imaging with sample rotation and multi-view fusion. Representative images of sagittal, axial and coronal views in original view of sample rotation (View A, left panel), 90° sample rotation (View B, middle panel) and fused image (right panel). Shown are xy, xz and yz sections from View A, rotated View B, and the fused reconstruction. Cartoons demonstrate sample orientation during both imaging runs.

b: Representative images of xy, yz and xz views in original view of sample rotation (View A, left panel), 90° sample rotation (View B, middle panel) and fused image (right panel). Scale bar = 20 µm.

c: Representative images of xy, yz and xz views in original view of sample rotation (View A, left panel), $90^{o}$ sample rotation (View B, middle panel) and fused image (right panel). Scale bar = 10 μm.

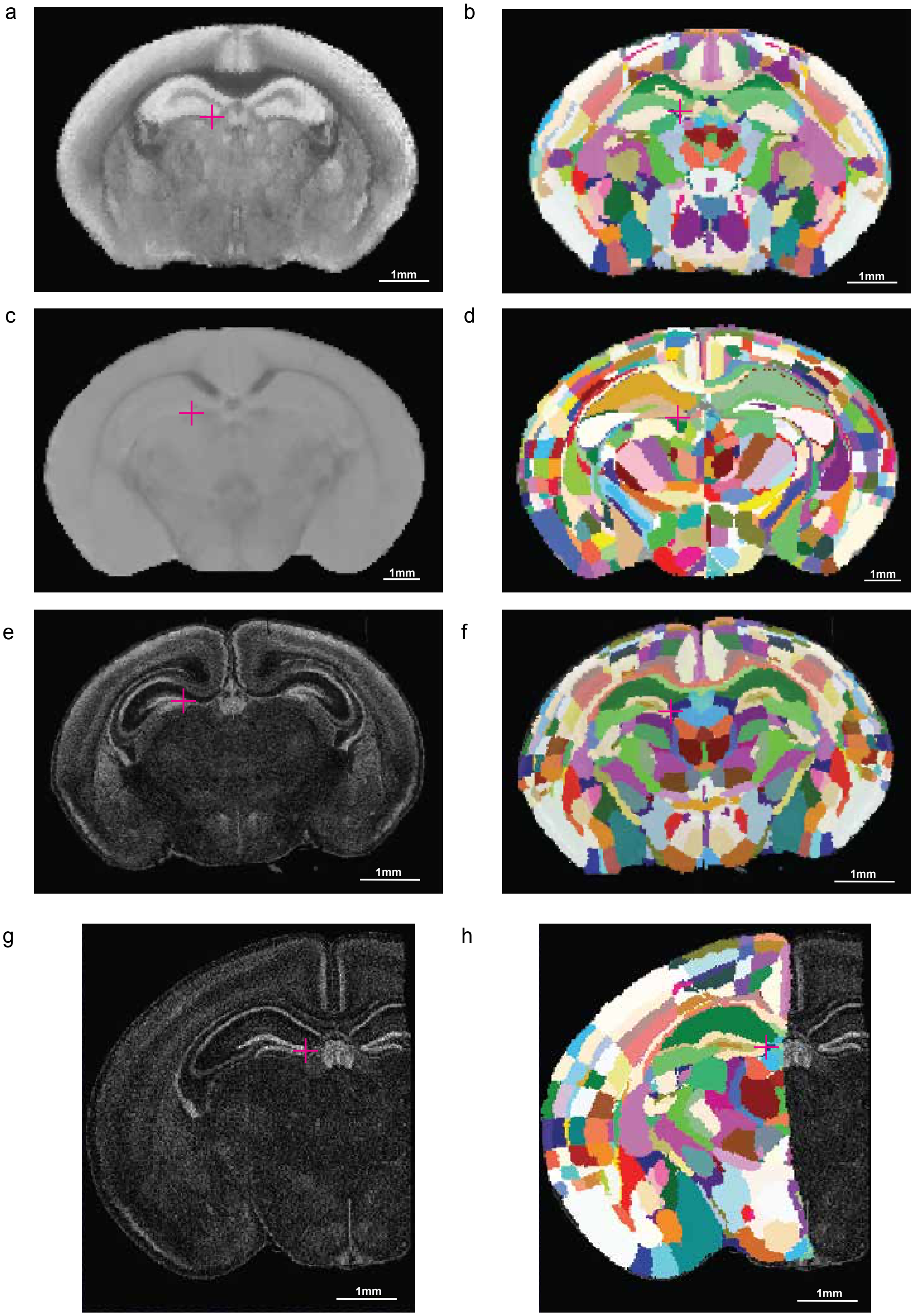


**Figure S4: MRI and Light-Sheet Registrations.**

a: Representative coronal MRI slice of P4 mouse brain. Purple (+) represents an anatomical location in MRI and the same region in (b) after co-registration with the atlas.

b: Corresponding P4 anatomical segmentation atlas overlaid on the MRI slice, illustrating region-specific annotations for early postnatal development. Each color represents a different annotated region from the atlas.

c: Representative coronal MRI slice of P14 mouse brain. Purple (+) represents an anatomical location in MRI and the same region in (d) after co-registration with the atlas. Contrast difference compared to (a) likely arises from applying the same MRI preparation protocol (including gadolinium concentration and incubation times) to brains of different sizes, but does not affect registration quality in (b) and (d).

d: Corresponding P14 anatomical segmentation atlas overlaid on the MRI slice.

e: High-resolution coronal section obtained via Light-Sheet Fluorescence Microscopy (LSFM) of a cleared P4 brain. Purple (+) represents an anatomical location in light-sheet image and the same region in (f) after co-registration with the atlas.

f: Corresponding atlas annotation mapped to the P4 LSFM data, showcasing the registration of 3D microscopy to a standard reference space.

g: Hemispheric coronal section of a cleared P14 brain using LSFM. Purple (+) represents an anatomical location in light-sheet image and the same region in (h) after co-registration with the atlas.

h: Corresponding atlas annotation mapped to the P14 LSFM data.

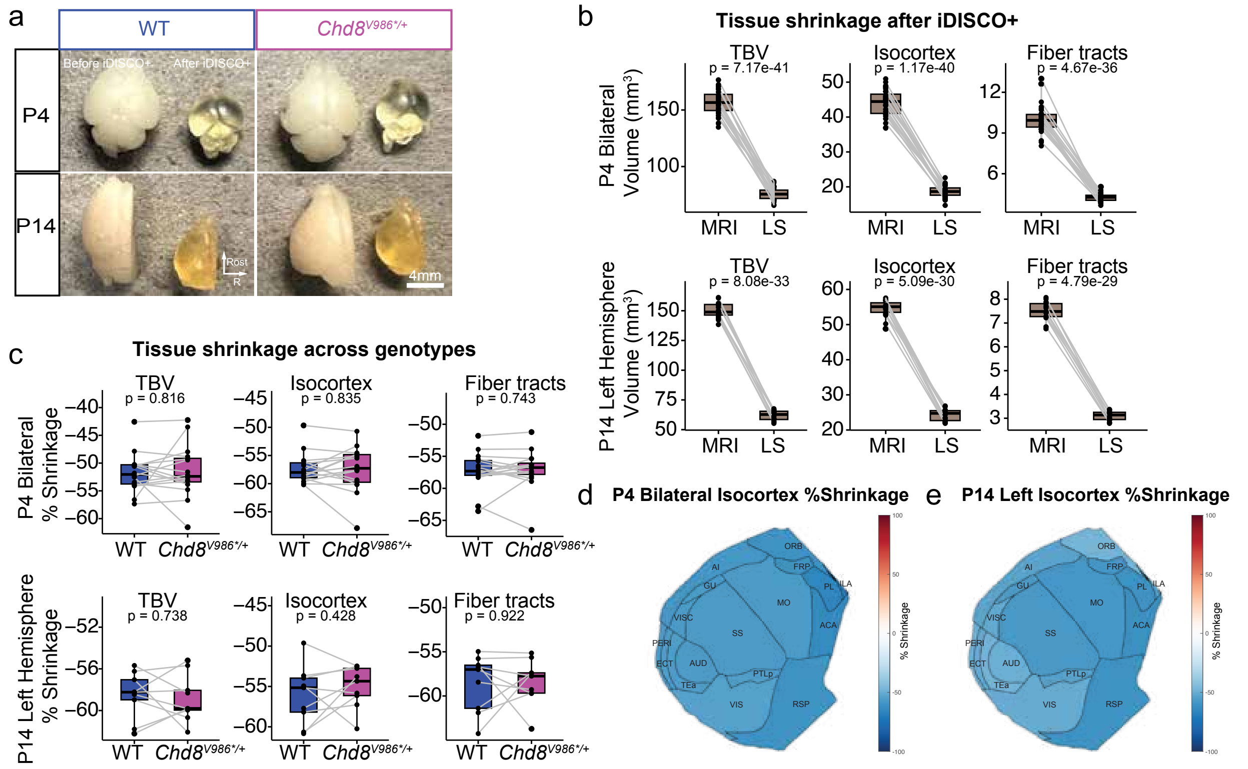


**Figure S5: Quantification of Tissue Shrinkage Following iDISCO+ Clearing.**

a: Representative macroscopic images of WT and *Chd8*$^{V986*/+}$ mouse brains at P4 and P14 before (left) and after tissue (right) clearing.

b: Paired plots showing the absolute volume (mm$^3$) of the total brain (TBV), isocortex and fiber tracts as measured by MRI (pre-clearing) versus light-sheet imaging (LS) (post-clearing) for P4 (top row; P-value is evaluated using a multiple linear regression controlling for sample, $n_{samples}$=30) and P14 (bottom row, P-value is evaluated using a multiple linear regression controlling for sample, $n_{samples}$=18).

c: Percentage shrinkage for TBV, isocortex and fiber tracts compared between WT and *Chd8*$^{V986*/+}$ mice for P4 (top row; P-value is evaluated using a multiple linear regression controlling for sample, $n_{samples}$=30) and P14 (bottom row, P-value is evaluated using a multiple linear regression controlling for sample, $n_{samples}$=18).

d: Flat-map projection of the P4 bilateral isocortex showing the percentage of shrinkage across functional cortical areas ($n_{samples}$=30).

e: Flat-map projection of the P14 left hemisphere isocortex showing the percentage of shrinkage across functional cortical areas ($n_{samples}$=18).

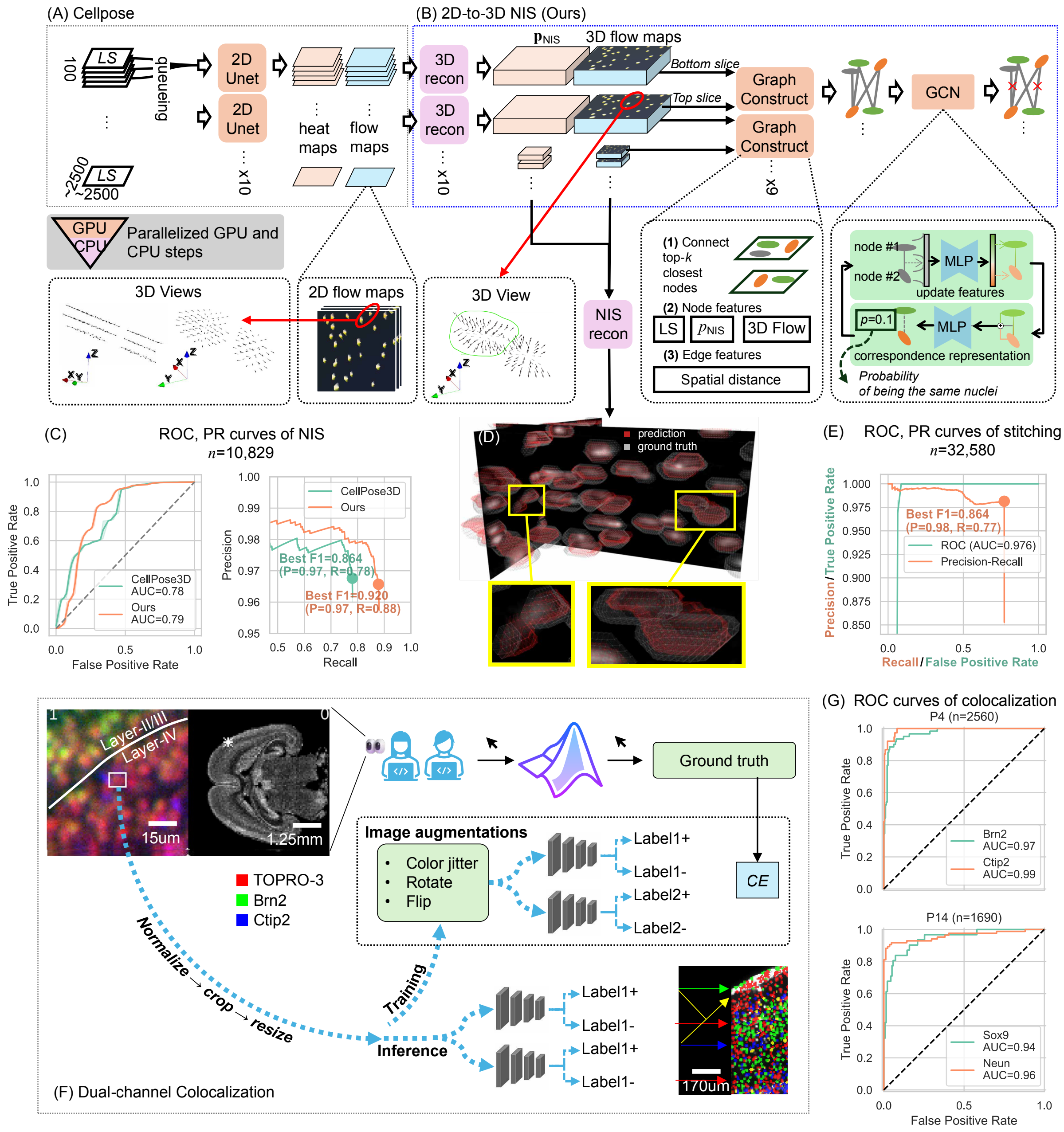

**Figure S6: Frameworks and Quantitative Accuracy of NIS and Colocalization.**

Our NIS framework is designed as a 2D-to-3D approach using (A) Cellpose backbone for 2D prediction, and (B) a graph neural network (GNN) based 3D reconstruction and stitching, where 2D Unet is built by a downsampling ResNet50 and an upsampling ResNet50, 3D reconstruction is the difference between gradient length of every 3rd and 1st slices after applying 7-layer median filter with kernel 3 and stride 1, and the NIS reconstruction from the flow and heat maps refers to Cellpose. (C) The ROC and Precision-Recall (PR) curves of NIS tested on n=10,829 3D annotations. (D) The visualization of our 3D NIS in one annotated volumetric image. (E) The ROC and PR curves of 3D NIS stitching tested on n=32,580 cropped NIS extracted from the 3D annotations. (F) The workflow of NIS colocalization: First, n=6,788 and n=3,889 for P4 and P14 annotations were manually collected using a Matlab GUI selectively picked from whole-brain NIS datasets among n=11 and n=6 P4 and P14 brains, respectively, by two neuroscience experts reviewing channel aligned three-channel image patches. The channel alignment is accomplished by Pearson Correlation Correction and a whole-brain density map is displayed aside to inform the global location of the patch. Then, each timepoint cohort has two ResNet50 models trained upon the manual annotations to predict if a channel (Brn2/Sox9 and Ctip2/Neun) is positive. Last, the entire whole-brain NIS results are iterated by the trained models to predict four classes of colocalization, as shown by red (both two channels are negative), green (Brn2+/Sox9+), blue (Ctip2+/Neun+), and yellow (both two channels are positive). (G) The ROC curves of colocalization by four models for Brn2, Ctip2, Sox9, and NeuN.

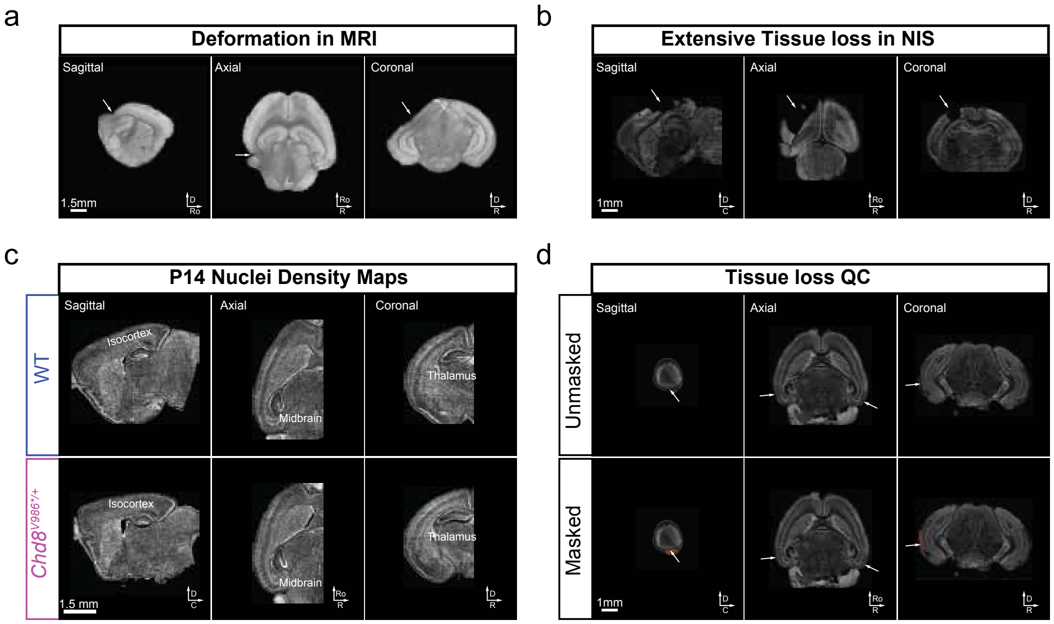


**Figure S7: Quality Control and Aberrations in Imaged Samples.**

a: Representative sagittal, axial, and coronal views of a magnetic resonance imaging (MRI) scan illustrating examples of tissue deformation that can occur during the preparation prior to imaging. Brains are stabilized in a syringe as previously described. However, the syringe piston may push into the brain to create deformities. Arrows point to specific areas where the brain tissue appears deformed.

b: Sagittal, axial, and coronal views of a brain imaged after clearing and nuclear staining (NIS), showing examples of significant tissue loss that can occur during sample preparation. Arrows indicate regions where tissue is missing or damaged. This sample was removed from analysis.

c: Representative sagittal, axial, and coronal nuclei density maps for Wild Type (WT, top row) and *Chd8*$^{V986*/+}$ (bottom row) mice at P14. Major brain regions, including the isocortex, midbrain, and thalamus, are labeled to show the distribution of nuclei across these structures.

d: Example of the quality control procedure for handling tissue loss. The top row ("Unmasked") shows raw images with arrows pointing to areas of tissue damage or loss. The bottom row ("Masked") shows the same images with these damaged regions highlighted in red, indicating they have been manually traced slice by slice to acquire the tissue loss volume and adjusted in a multiple linear regression controlling for pair and manually annotated tissue loss: Phenotype ~ Genotype + Pair + Tissue loss.

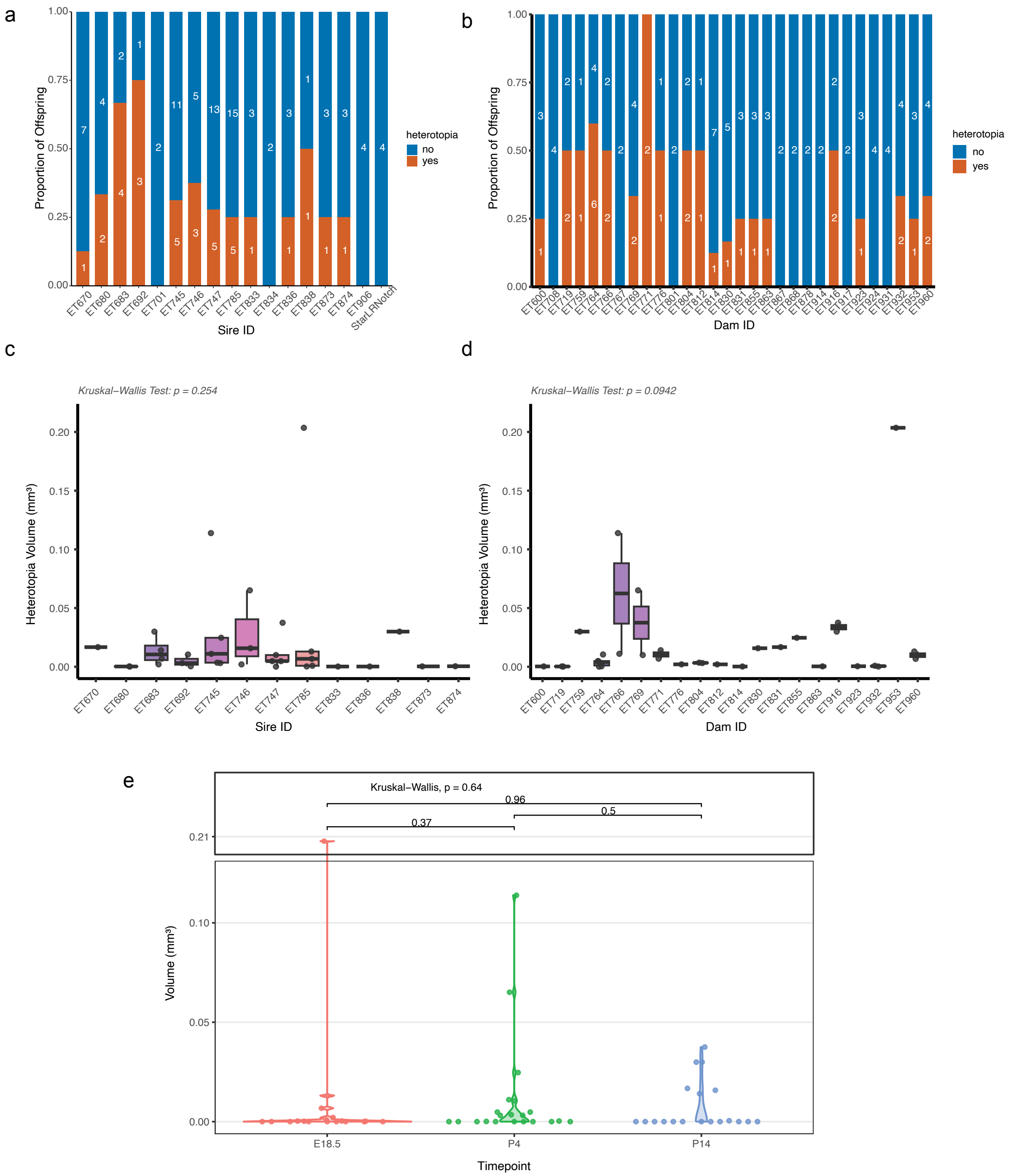


**Figure S8: Heterotopia Incidence and Volume across Sires, Dams and Developmental Timepoints.**

a & b: Stacked bar charts showing the proportion of offspring with (orange) and without (blue) heterotopia, categorized by **(a)** Sire ID (all *Chd8*$^{V986*/+}$) and **(b)** Dam ID (all WT). Numbers within bars indicate the total count of offspring per parent. Heterotopias were present in litters from multiple dams and sires, so were not likely caused by ungenotyped mutations in one breeding pair.

c & d: Box-and-whisker plots overlaid with individual data points representing the total heterotopia volume ($mm^3$) per affected offspring, grouped by **(c)** Sire ID and **(d)** Dam ID. Differences were evaluated using Kruskal-Wallis tests. No significant differences were found indicating one breeding pair is not driving heterotopia size.

e: Violin plots showing the distribution of heterotopia volumes at three key timepoints: E18.5, P4, P14. Differences were evaluated using Kruskal-Wallis tests. No significant differences were found indicating heterotopias have similar volumes through development.

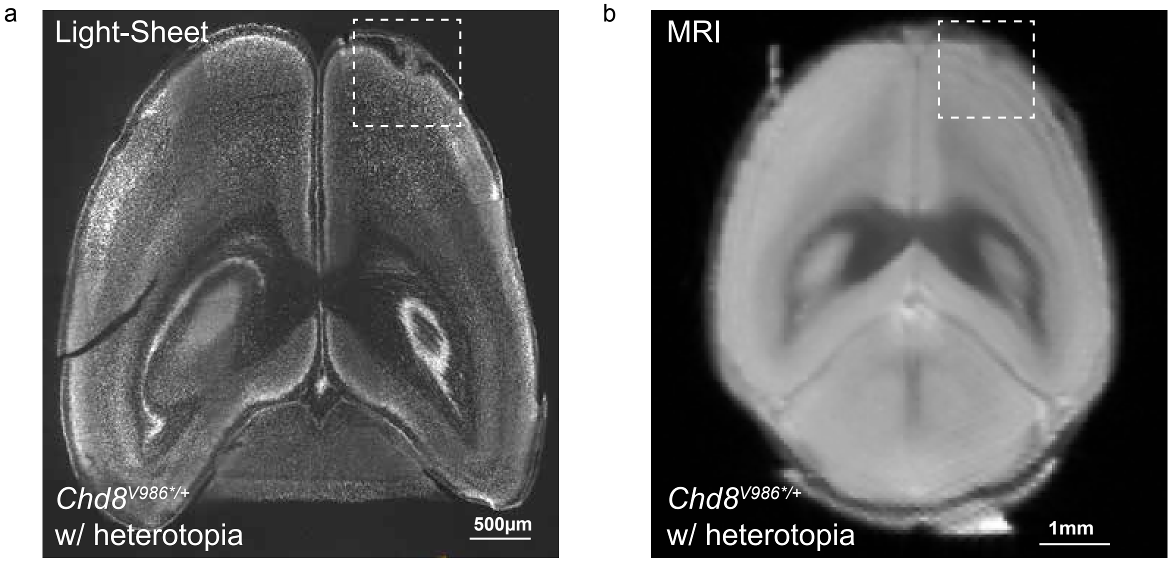


**Figure S9: Visualization of Molecular Layer Heterotopia in *Chd8*$^{V986*/+}$ Mice using Light-Sheet Microscopy and MRI.**

a: Representative Light-Sheet fluorescence microscopy image of a coronal section from a *Chd8*$^{V986*/+}$ mouse brain. A white dashed box highlights a region of cortical malformation (heterotopia) in the upper right hemisphere, revealing high-resolution structural details of the lesion.

b: Corresponding coronal MRI slice of a *Chd8*$^{V986*/+}$ mouse brain. The white dashed box indicates the same anatomical region as in (a), demonstrating that the heterotopia is detectable as a subtle gross morphological anomaly even at the lower resolution of MRI.

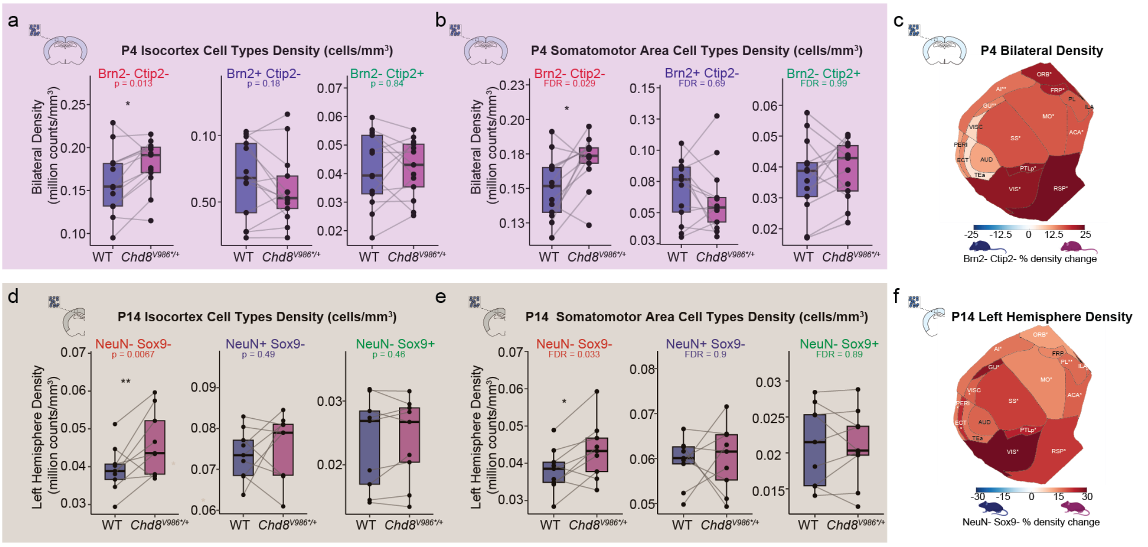


**Figure S10: Early Density Increases at P4 in *Chd8*$^{V986*/+}$ Brain Precedes Brain Volume Expansion at P14.**

a: Cell density (million counts/mm$^3$) for the three classified populations across P4 bilateral isocortex ($n_{pairs}$ = 13). Lines indicate littermate pairs with P-value evaluated using a multiple linear regression controlling for pair and tissue loss.

b: Cellular density (million counts/mm$^3$) within the P4 bilateral somatomotor area for the three classified populations ($n_{pairs}$ = 13). Lines indicate littermate pairs with p-value evaluated using a multiple linear regression controlling for pair and tissue loss. Significance is indicated by False Discovery Rate (* FDR < 0.1, ** FDR < 0.01, *** FDR < 0.001).

c: Flattened isocortex heatmap of P4 Brn2-/Ctip2- bilateral cell density changes displaying the percentage change in Brn2-/Ctip2- density between WT and *Chd8*$^{V986*/+}$ littermate pairs across 17 isocortical regions ($n_{pairs}$ = 13). P-value evaluated using a multiple linear regression controlling for pair and tissue loss. Significance is indicated by False Discovery Rate (* FDR < 0.1, ** FDR < 0.01, *** FDR < 0.001).

d: Cell density (million counts/mm$^3$) for the three classified populations across P14 left hemisphere isocortex ($n_{pairs}$ = 9). Lines indicate littermate pairs with P-value evaluated using a multiple linear regression controlling for pair and tissue loss.

e: Cellular density (million counts/mm$^3$) within the P14 left hemisphere somatomotor area for the three classified populations ($n_{pairs}$ = 9). Lines indicate littermate pairs with p-value evaluated using a multiple linear regression controlling for pair and tissue loss. Significance is indicated by False Discovery Rate (* FDR < 0.1, ** FDR < 0.01, *** FDR < 0.001).

f: Flattened isocortex heatmap of P14 NeuN-/Sox9- left hemisphere cell density changes displaying the percentage change in NeuN-/Sox9- density between WT and *Chd8*$^{V986*/+}$ littermate pairs across 17 isocortical regions ($n_{pairs} = 9$). P-value evaluated using a multiple linear regression controlling for pair and tissue loss. Significance is indicated by False Discovery Rate (* FDR < 0.1, ** FDR < 0.01, *** FDR < 0.001).

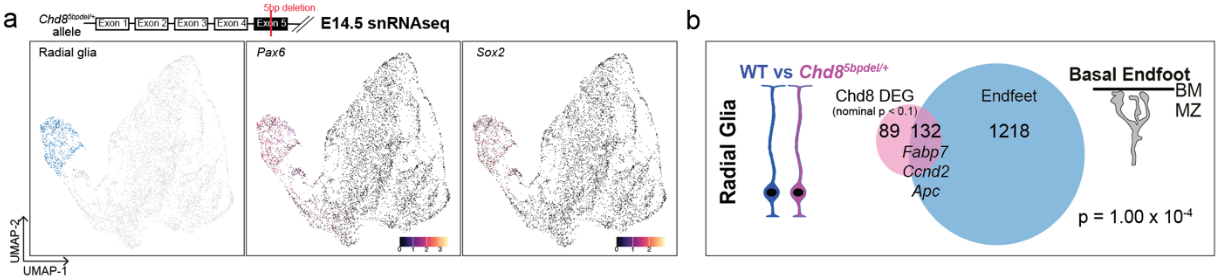


**Figure S11: Validation of Radial Glial Endfoot Transcriptomic Dysregulation in an Independent *Chd8* Mutant Mouse Line.**

a: Schematic of the *Chd8*$^{5bpdel/+}$ allele targeting Exon 5 (top) and UMAP visualization of E14.5 single-nucleus RNA-sequencing (snRNA-seq) data (bottom). The blue cluster highlights the localized radial glial cell population (795 cells, n = 4), which is characterized by the robust expression of canonical lineage markers *Pax6* and *Sox2*.

b: Venn diagram illustrating a significant transcriptomic overlap ($p = 1.00 \times 10^{-4}$) between nominal *Chd8* single-nucleus differentially expressed genes (DEGs, nominal $p < 0.1$; n = 221) and a physically isolated cortical basal endfoot gene profile (n = 1350). The intersection identifies 132 shared targets, where several highlighted downstream *Chd8* target genes, including *Fabp7*, *Ccnd2*, and *Apc*, are shared across models (Figure 7k). BM, basement membrane; MZ, marginal zone.

**Supplemental Table legends:**

**Table S1:** Summary of E18.5, P4 and P14 samples used for each analyses across multiple imaging modalities.

**Table S2:** Summary statistics of volumetric analyses across cortical regions between WT and *Chd8*$^{V986*/+}$ at P4 (sheet 1) and P14 (sheet 2) in $mm^3$.

**Table S3:** Summary statistics of nuclei segmentations across cortical regions between WT and *Chd8*$^{V986*/+}$ at P4 (sheet 1) and P14 (sheet 2) in million count.

**Table S4:** Summary statistics of density analyses across cortical regions between WT and *Chd8*$^{V986*/+}$ at P4 (sheet 1) and P14( sheet 2) in million counts/$mm^3$.

**Table S5:** Summary statistics of Brn2-Ctip2- (sheet 1), Brn2-Ctip2+ (sheet 2), Brn2+Ctip2- (sheet 3) for P4 timepoint and NeuN-Sox9- (sheet 4), NeuN-Sox9+ (sheet 5) and NeuN+Sox9- (sheet 6) for P14, colocalization with TO-PRO-3 across cortical regions between WT and *Chd8*$^{V986*/+}$ in million count.

**Table S6:** Summary statistics of Brn2-Ctip2- (sheet 1), Brn2-Ctip2+ (sheet 2), Brn2+Ctip2- (sheet 3) for P4 timepoint and NeuN-Sox9- (sheet 4), NeuN-Sox9+ (sheet 5) and NeuN+Sox9- (sheet 6) for P14, colocalization with TO-PRO-3 across cortical regions between WT and *Chd8*$^{V986*/+}$ million counts/$mm^3$.

**Table S7:** Raw values of MRI, NIS, Tissue loss and Density for WT and *Chd8*$^{V986*/+}$ samples at P4.

**Table S8:** Raw values of MRI, NIS, Tissue loss and Density for WT and *Chd8*$^{V986*/+}$ samples at P14.

**Table S9:** Raw values of Brn2-Ctip2- (sheet 1), Brn2+Ctip2- (sheet 2), Brn2-Ctip2+ (sheet 3) for P4 timepoint and NeuN-Sox9- (sheet 4), NeuN+Sox9- (sheet 5) and NeuN-Sox9+ (sheet 6) at P14, colocalization with TO-PRO-3 across cortical regions for all samples at P14.

**Table S10:** Raw values of Brn2-Ctip2- (sheet 1), Brn2+Ctip2- (sheet 2), Brn2-Ctip2+ (sheet 3) for P4 timepoint and NeuN-Sox9- (sheet 4), NeuN+Sox9- (sheet 5) and NeuN-Sox9+ (sheet 6) at P14, colocalization Density across cortical regions for all samples at P14.

**Table S11:** Raw values of tissue shrinkage and summary statistics for WT and *Chd8*$^{V986*/+}$ mice at P4 (sheet 1 and 2) and P14 (sheet 3 and 4).

**Table S12:** Number of training data for NIS (sheet 1) and colocalization (sheet 2) organized by sex, genotype and time point.

**Supplementary Video Legends:**

**Video S1: Representative Images of all markers of WT and *Chd8*$^{V986*/+}$ brains at P4.**

Immunofluorescent markers for of all markers of WT and *Chd8*$^{V986*/+}$ brains at P4 in 2D and 3D views. TO-PRO-3 = grey, Brn2 = red, Ctip2 = blue.

**Video S2: Representative Images of all markers of WT and *Chd8*$^{V986*/+}$ brains at P14.**

Immunofluorescent markers for of all markers of WT and *Chd8*$^{V986*/+}$ brains at P14 in 2D and 3D views. TO-PRO-3 = grey, NeuN = cyan, Sox9 = magenta

**Video S3: 3D Volumetric Reconstruction and 2D Cross-Sectional Views of Molecular Layer Heterotopia in E18.5 *Chd8*$^{V986*/+}$ brain.**

A comprehensive spatial visualization of a representative molecular layer heterotopia in 3D and 2D in a E18.5 *Chd8*$^{V986*/+}$ mouse. TO-PRO-3 = grey.

**Video S4: 3D Volumetric Reconstruction and 2D Cross-Sectional Views of Molecular Layer Heterotopia in P4 *Chd8*$^{V986*/+}$ brain.**

A comprehensive spatial visualization of a representative molecular layer heterotopia in 3D and 2D in a P4 *Chd8*$^{V986*/+}$ mouse. TO-PRO-3 = grey, Brn2 = green, Ctip2 = magenta.

**Video S5: 3D Volumetric Reconstruction and 2D Cross-Sectional Views of Molecular Layer Heterotopia in P14 *Chd8*$^{V986*/+}$ brain.**

A comprehensive spatial visualization of a representative molecular layer heterotopia in 3D and 2D in a P14 *Chd8*$^{V986*/+}$ mouse. NeuN = cyan.
